# Pixel2Gene enables histology-guided reconstruction and prediction of spatial gene expression

**DOI:** 10.64898/2026.02.21.707168

**Authors:** Sicong Yao, Amelia Schroeder, Shunzhou Jiang, Soyoung Im, Jeong Hwan Park, Bernhard Dumoulin, Tae Hyun Hwang, Katalin Susztak, Mingyao Li

**Affiliations:** Statistical Center for Single-Cell and Spatial Genomics, Department of Biostatistics, Epidemiology and Informatics, University of Pennsylvania Perelman School of Medicine, Philadelphia, PA, USA; Department of Computational Biology, St. Jude Children’s Research Hospital, Memphis, TN, USA; Department of Pathology, St. Vincent’s Hospital, College of Medicine, The Catholic University of Korea, Seoul, Republic of Korea; Department of Pathology, Seoul Metropolitan Government-Seoul National University Baramae Medical Center, Seoul National University College of Medicine, Seoul, Republic of Korea; Renal, Electrolyte, and Hypertension Division, Department of Medicine, University of Pennsylvania Perelman School of Medicine, Philadelphia, PA, USA; Penn/CHOP Kidney Innovation Center, University of Pennsylvania Perelman School of Medicine, Philadelphia, PA, USA; Department of Surgery, Vanderbilt University Medical Center, Nashville, TN, USA; Center for Systems Biology, Vanderbilt University, Nashville, TN, USA; Department of Pathology and Laboratory Medicine, University of Pennsylvania Perelman School of Medicine, Philadelphia, PA, USA

## Abstract

Advances in spatial transcriptomics (ST) have fundamentally transformed our understanding of tissue biology by enabling gene expression profiling within intact spatial contexts and uncovering tissue organization and microenvironmental interactions. However, current high-resolution ST platforms remain constrained by high costs, limited tissue coverage, and technical artifacts, often yielding noisy, sparse, and incomplete data that compromise analytical accuracy, biological interpretation, and clinical utility. To address these challenges, we introduce Pixel2Gene, a deep learning framework that integrates co-registered histology images with ST data to enable histology-guided reconstruction and prediction of spatial gene expression. Pixel2Gene enhances existing expression measurements by denoising low-confidence data and reconstructing coherent expression patterns, while also predicting gene expression in unmeasured tissue regions and new samples lacking direct transcriptomic profiling. We systematically evaluated Pixel2Gene across multiple high-resolution ST platforms, including Visium HD, Xenium, and CosMx, spanning diverse tissue types and disease contexts using downsampling simulations and cross-platform comparisons in clinical samples. Across all settings, Pixel2Gene consistently improved data consistency, mitigated dropout effects, restored biologically meaningful spatial structure, and enabled accurate downstream analyses. By leveraging the scalability and ubiquity of routine histology, Pixel2Gene facilitates comprehensive, cost-effective ST profiling at whole-tissue scale, supporting large cohort studies, translational research, and next-generation biomarker discovery.

## Introduction

Spatial transcriptomics (ST) enables gene expression profiling within intact tissues, offering a powerful framework to investigate cellular organization, molecular heterogeneity, and cell-cell interactions *in situ*^1,2^. Unlike dissociative single-cell sequencing methods, ST preserves tissue architecture, providing critical insights into development, disease progression, and immune dynamics^3^. The field has rapidly evolved from early spot-based platforms with multicellular resolution to high-resolution technologies such as NanoString’s CosMx^4^, Vizgen’s MERSCOPE^5^, and 10x Genomics’ Visium HD^6^ and Xenium^7^, which achieve near-cellular or even subcellular resolution. These next-generation platforms enable fine-grained single-cell mapping, detection of rare cell populations, and characterization of spatial gradients across diverse tissue types, substantially expanding the analytical scope of spatial omics.

Despite these advances, high-resolution ST platforms still face significant limitations. High costs, lengthy processing times, and restricted fields of view (FOVs) limit scalability and prevent comprehensive tissue analysis^8,9^. Even within profiled regions, incomplete capture of molecules results in sparsity and technical artifacts, such as dropout and signal leakage. For instance, although Visium HD enables gene expression profiling at 2 µm, 8 µm, and 16 µm bin sizes, the resulting expression maps are often patchy and discontinuous, which complicates downstream analyses that require coherent spatial patterns. Similarly, CosMx generates sparse and fragmented tissue coverage across discrete FOVs, which limits comprehensive tissue characterization. These challenges hinder the robust analysis of heterogeneous or large specimens, where complete and reliable gene expression maps are most crucial.

Nevertheless, high-resolution ST data do retain substantial biological signals, where many genes show consistent and biologically meaningful expression patterns in certain regions, even when others are degraded or sparsely captured. This observation presents an opportunity for computational methods to leverage high-confidence regions and underlying tissue structure to recover, enhance, and predict gene expression in low-quality or unmeasured tissue areas, thereby extending the utility of existing ST datasets beyond their experimental limitations.

Hematoxylin and eosin (H&E) staining provides a powerful complement to ST. Routinely used in clinical practice, H&E captures nuclear morphology, cellular organization, and tissue architecture, representing features that reflect underlying gene expression states. Whole-slide imaging achieves subcellular resolution (<0.5 µm/pixel)^10^ across centimeter-scale tissue sections, with rapid turnaround (<24 hours)^11^ and minimal cost (∼$10 per slide)^12^, making it both high-quality and highly scalable. Prior studies have demonstrated strong correlations between histological features and transcriptomic profiles, suggesting that histomorphology can serve as a surrogate for molecular state^13–22^. These observations raise an opportunity: by systematically learning the relationship between histological features and reliable gene expression in high-quality ST regions, one can enhance gene expression quality in low-quality regions and computationally predict gene expression where direct measurements are missing or unreliable. Integrating histology with ST thus leverages the complementary strengths of both modalities and enables the reconstruction and prediction of accurate, high-resolution molecular tissue maps that overcome the limitations of current ST platforms.

In this paper, we present Pixel2Gene, a deep learning framework that enhances single-cell resolution ST data by integrating co-registered histology images. Unlike prior methods^23,24^ that aim solely to infer gene expression de novo from histology, Pixel2Gene jointly leverages high-confidence ST measurements and histological features to both enhance existing expression data and predict gene expression in regions with low-quality or missing measurements. By integrating preprocessed high-quality transcriptomic data with histological features in a unified framework, Pixel2Gene reconstructs biologically faithful expression maps that scale to whole-slide tissue sections while preserving native architecture. Our results demonstrate that Pixel2Gene maintains genuine biological signals, mitigates dropout and platform-specific artifacts, and enables accurate, biologically meaningful spatial analyses that uncover fine-grained tissue organization and disease-associated patterns.

## Results

### Overview of Pixel2Gene

An overview of the Pixel2Gene workflow is shown in **Fig. 1**. The framework aims to reconstruct high-resolution spatial gene expression maps by integrating co-registered H&E images with filtered single-cell resolution ST data, to enhance existing measurements and infer expression in unmeasured tissue regions. The pipeline begins by aligning the two modalities within a shared coordinate system. Here, 16 × 16-pixel H&E superpixels, approximately 8 µm × 8 µm in size, are paired with gene expression data assigned to the corresponding superpixels, either directly from 8 µm ST bins in Visium HD or by aggregating molecule counts according to transcript coordinates. A tissue mask is then derived from HistoSweep^25^, a high-resolution H&E image quality control and preprocessing method to identify biologically relevant regions and exclude artifacts such as tissue folds or staining debris, ensuring that analysis focuses on informative tissue areas.

**Fig. 1.**
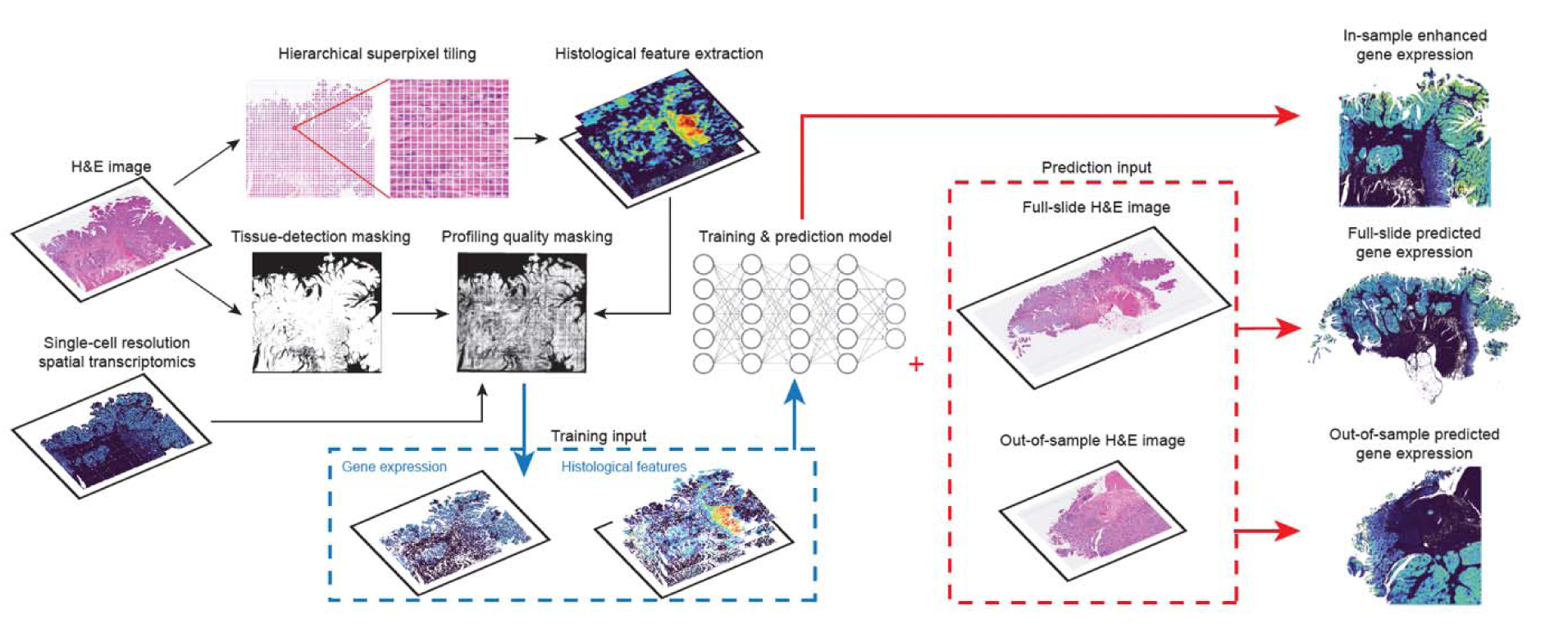
Overview of Pixel2Gene workflow. Pixel2Gene reconstructs high-resolution spatial gene expression maps by integrating co-registered H&E images with filtered single-cell resolution ST data. H&E images and ST measurements are spatially aligned into a shared coordinate framework, where histological features are extracted hierarchically at superpixel resolution. Pixel2Gene applies semantic tissue masks to exclude artifacts and aggregates transcriptomic signals into spatially coherent clusters. Within each cluster, high-confidence ST data guide model training to learn histology-expression relationships using a feed-forward neural network. Pixel2Gene enhances noisy gene expression within profiled regions by leveraging spatial and morphological context to recover biologically coherent patterns. Beyond in-tissue enhancement, it also enables full-tissue reconstruction and out-of-sample prediction—inferring gene expression in unprofiled areas or entirely new tissue sections using only H&E images.

Next, Pixel2Gene extracts multiscale histological features for each superpixel, capturing nuclear morphology, local cellular neighborhoods, and broader microenvironmental context. These features are used to segment the tissue into histologically coherent regions, which guide quality filtering and localized model training. Within each cluster, superpixels with UMI counts below a predefined threshold are excluded to retain only high-quality data for robust model learning.

Using these filtered data pairs, a feed-forward neural network is trained to capture the relationship between histology and gene expression. The resulting model serves two complementary functions: (1) it enhances existing ST by denoising low-confidence measurements, and (2) it infers gene expression *de novo* in unmeasured regions, including gaps within the ST field of view or areas lacking transcriptomic profiles. This integrated framework generates enhanced, biologically faithful expression maps that preserve native tissue architecture and scale to whole-slide specimens, enabling practical analysis of datasets containing hundreds of thousands to millions of spatial units (**Supplementary Tables 1 and 2**). These reconstructed maps enable downstream applications such as cell type annotation, spatial domain detection, and discovery of disease-associated tissue patterns at single-cell resolution.

### Downsampling evaluation on human colorectal cancer and mouse brain samples

To rigorously assess Pixel2Gene’s ability to recover accurate gene expression from degraded ST data, we performed downsampling simulations on two high-resolution datasets: a human colorectal cancer sample (CRC-P2) from patient 2 and a fresh-frozen mouse brain sample, both profiled using 10x Genomics Visium HD. These datasets provided dense, spatially resolved gene expression profiles paired with co-registered H&E images, providing an ideal testbed for simulating realistic data sparsity while maintaining biological relevance.

Since ground truth expression values are not directly available due to the inherent noise and incompleteness of single-cell resolution ST data, we restricted our evaluation to the top 300 highly expressed genes, which are less susceptible to dropout and provide a more stable reference for assessing reconstruction accuracy. To benchmark Pixel2Gene’s performance, we simulated degraded input data by applying a controlled downsampling procedure that mimics spatial variability and technical sparsity observed in low-quality ST experiments (**Methods**).

Pixel2Gene was trained to reconstruct high-fidelity gene expression maps using downsampled ST data and the corresponding H&E images in CRC-P2 as input. Comparative analysis revealed Pixel2Gene’s superior ability to recover coherent spatial patterns from degraded inputs while preserving critical tissue structures. We highlight five representative genes, including *KRT8*, *SPARC*, *PLAC8*, *SELENOP*, and *TAGLN*, selected for their distinct spatial expression profiles and tissue-specific distributions (**Fig. 2a**). Pixel2Gene accurately recovered their characteristic expression patterns, preserving expression gradients and anatomical boundaries that were obscured in the downsampled data.

**Fig. 2.**
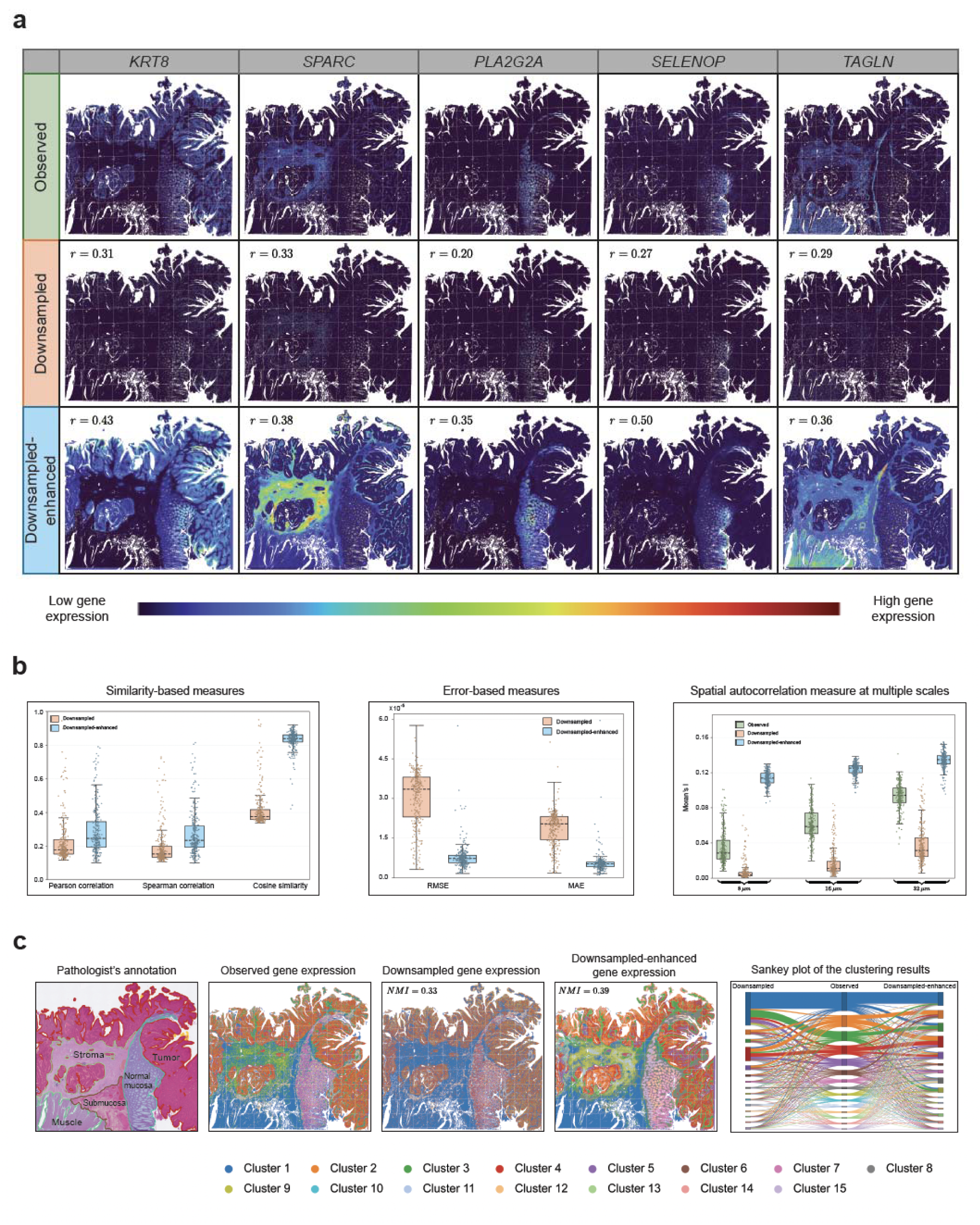
Evaluation of Pixel2Gene enhancement in the CRC simulation data. **a**, Visualization of spatial expression patterns of selected genes in CRC-P2 Visium HD data at 8 µm resolution, shown for the observed, downsampled, and Pixel2Gene-enhanced (downsampled-enhanced) gene expression. **b**, Quantitative comparison of prediction accuracy using Pearson correlation, Spearman correlation, cosine similarity (higher is better), RMSE, and MAE (lower is better), together with spatial autocorrelation measured by Moran’s I (higher is better), computed across the top 300 highly expressed genes. Pearson, Spearman, Cosine, RMSE, and MAE were calculated by comparing observed expression with downsampled or downsampled-enhanced predictions at 8 µm resolution. Moran’s I was computed independently for each modality across multiple spatial resolutions. In each boxplot, boxes indicate the interquartile range (IQR; 25th–75th percentile), the center dashed line denotes the median, and whiskers extend to the most extreme values within 1.5×IQR **c**, Comparison of spatial clustering results across different input modalities using top 300 highly expressed genes. From left to right: pathologist annotation (serving as biological reference), clustering based on observed gene expression, downsampled gene expression, and Pixel2Gene-enhanced expression. Cluster labels for the downsampled and enhanced results were aligned to the observed clustering using a one-to-one assignment (Hungarian algorithm) based on the overlap of spot assignments, enabling consistent color coding across panels. NMI scores with respect to the observed clustering are shown in the top-left corner of each panel. The Sankey diagram summarizes the correspondence of cluster memberships across downsampled (left), observed (middle), and enhanced (right) results after label alignment.

To quantitatively evaluate Pixel2Gene’s reconstruction accuracy, we compared the predicted expression values with the original expression values using five complementary metrics: Pearson correlation, Spearman correlation, cosine similarity, root mean squared error (RMSE), and mean absolute error (MAE). Across all metrics, Pixel2Gene consistently outperformed the downsampled inputs, demonstrating robust recovery of biological signal from degraded data **(Fig. 2b)**. Spatial autocorrelation analysis via Moran’s *I* at multiple scales (16 µm, 32 µm and 64 µm) further showed that Pixel2Gene not only recovered, but in some cases enhanced, the spatial organization relative to both the downsampled and the original data. These results validate the model’s dual capability to: (1) denoise technical artifacts, and (2) reinforce genuine biological patterns through histology-guided reconstruction.

Our unsupervised clustering analysis also highlighted the benefits of Pixel2Gene reconstruction. Clusters derived from the downsampled data failed to capture coherent tissue organization or biologically meaningful spatial domains. In contrast, clustering based on Pixel2Gene-enhanced expression successfully recovered fine-grained spatial patterns, in some regions even more distinctly than those observed in the original data, suggesting improved signal-to-noise ratios and enhanced resolution of tissue architecture (**Fig. 2c**). To quantitatively assess the biological coherence of reconstructed expression profiles, we computed the intra-cluster correlations (ICC) for observed, downsampled, and downsampled-enhanced data, using the unsupervised clustering labels derived from each respective dataset (**Supplementary Fig. 2a**). Pixel2Gene substantially improved ICC relative to the downsampled baseline and, in many cases, even exceeded the ICC of the original data, indicating that prediction not only restored but also enhanced the internal consistency of gene expression within biologically defined regions. Additionally, the normalized mutual information (NMI) between Pixel2Gene-enhanced and observed clusters was higher than that between downsampled and observed clusters, further supporting the biological consistency of the enhanced expression landscape.

We observed similar performance trends in the fresh-frozen mouse brain dataset, supporting the generalizability of Pixel2Gene across tissue types (**Supplementary Figs. 1 and 2b**). These findings demonstrate that Pixel2Gene effectively denoises and reconstructs sparse ST data by leveraging histological context and spatial structure, enabling high-fidelity recovery of gene expression across tissue sections, particularly for biologically important genes with spatial patterns.

### Pixel2Gene enables scalable enhancement across diverse ST platforms

While our downsampling experiments focused on the top 300 highly expressed genes as reliable benchmarks, standard Visium HD profiling captures the whole transcriptome, where the majority of genes are affected by substantial technical noise that hampers immediate biological interpretation. To evaluate Pixel2Gene’s ability to enhance transcriptome-wide data, we trained the model on a Visium HD sample derived from the same tissue block as the CRC-P2 Xenium dataset. Following Pixel2Gene reconstruction, the spatial expression profiles of those lower-confidence genes became markedly clearer and more biologically coherent, revealing spatial structure that was largely obscured in the original data (**Fig. 3**).

**Fig. 3.**
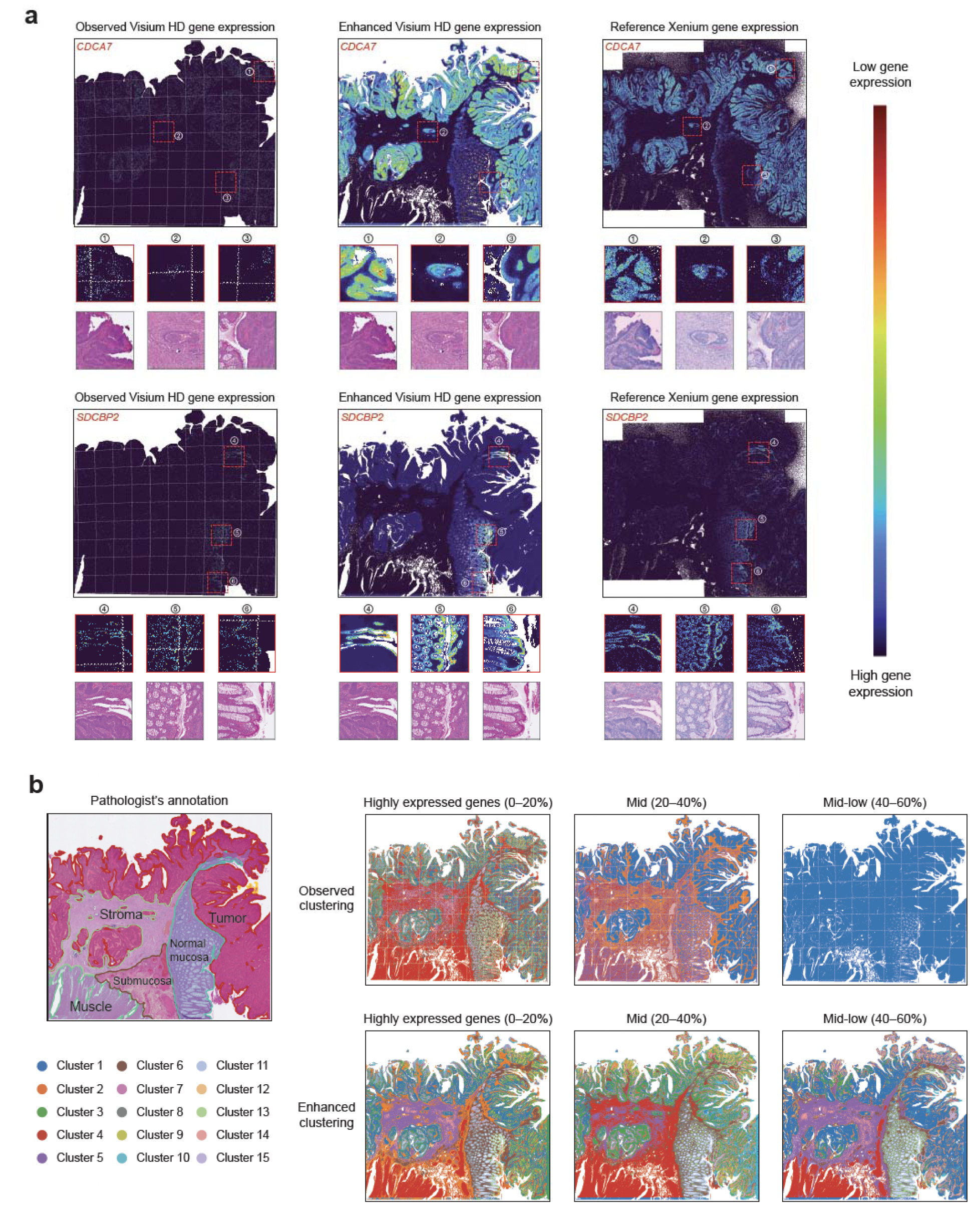
Pixel2Gene enhances transcriptome-wide expression and recovers spatial structure from noisy Visium HD data. **a**, Visualization of selected genes from Visium HD CRC-P2 not included in the top 300 highly expressed gene set. Expression of *CDCA7* and *SDCBP2* is shown for observed Visium HD data, Pixel2Gene-enhanced predictions, and matched adjacent Xenium measurements. Enhanced predictions yield sharper, spatially coherent patterns that align closely with high-resolution Xenium references, despite noise in the original Visium HD data. **b**, Unsupervised clustering using different expression abundance bins: top 0–20% (high), 20–40% (mid), and 40–60% (mid-low) highly expressed genes. Pixel2Gene-enhanced data reveals fine-grained spatial domains across all tiers, whereas clustering on observed expression fails to resolve tissue architecture, particularly for mid and mid-low expression bins. This demonstrates Pixel2Gene’s ability to recover biologically meaningful spatial organization even from weak expression signals.

As an illustrative example, *CDCA7*, a gene associated with mitotic epithelial cells, appeared sparse and fragmented in the raw Visium HD data, especially in the fine-grained tumor regions (**Fig. 3a**). After Pixel2Gene enhancement, its spatial pattern became markedly clearer and more coherent, closely aligning with expression in the adjacent high-resolution Xenium CRC-P2 dataset, which was used here as an independent reference. Similarly, *SDCBP2*, another gene outside the highly expressed gene set, exhibited enhanced spatial pattern after Pixel2Gene’s enhancement. These examples demonstrate Pixel2Gene’s ability to recover biologically spatial patterns even for noisy, low-confidence genes, yielding expression profiles comparable to those in the higher-quality Xenium CRC-P2 data. Additional examples further demonstrate that Pixel2Gene consistently recovers biologically coherent spatial patterns across genes with diverse and distinct spatial distributions (**Supplementary Fig. 3**).

To further evaluate biological utility beyond individual genes, we performed unsupervised clustering using genes stratified by expression level. For the top 0–20% highly expressed genes, Pixel2Gene enhancement revealed well-structured tissue compartments that were partially resolved in the observed data. Notably, the advantages of Pixel2Gene became especially pronounced in the mid-expression bins (20–40% and 40–60%), where native expression was sparse and noisy. In these tiers, typically enriched for functionally relevant but weakly expressed genes, clustering based on raw expression failed to capture coherent spatial domains. In contrast, Pixel2Gene recovered well-organized, biologically meaningful structures, uncovering tissue compartments and microenvironments that were otherwise obscured (**Fig. 3b**). These results demonstrate that Pixel2Gene not only enhances individual gene profiles but also uniquely restores latent spatial architecture across the entire expression spectrum, including mid-abundance genes that are essential for nuanced tissue characterization but often lost in conventional analyses.

We next evaluated Pixel2Gene’s generalizability on Xenium, an imaging-based spatial transcriptomics platform with higher resolution but lower gene coverage (∼400 genes). Applying Pixel2Gene to a centimeter-scale large gastric cancer tissue section profiled with 10x Xenium (DOI: 10.5281/zenodo.15164980)^26^, we observed marked improvements in spatial coherence across key marker genes, including *ACTA2*, *CD3D*, EPCAM, and *KRT20*, whose native Xenium expression was noisy or spatially fragmented (**Fig. 4a**).

**Fig. 4.**
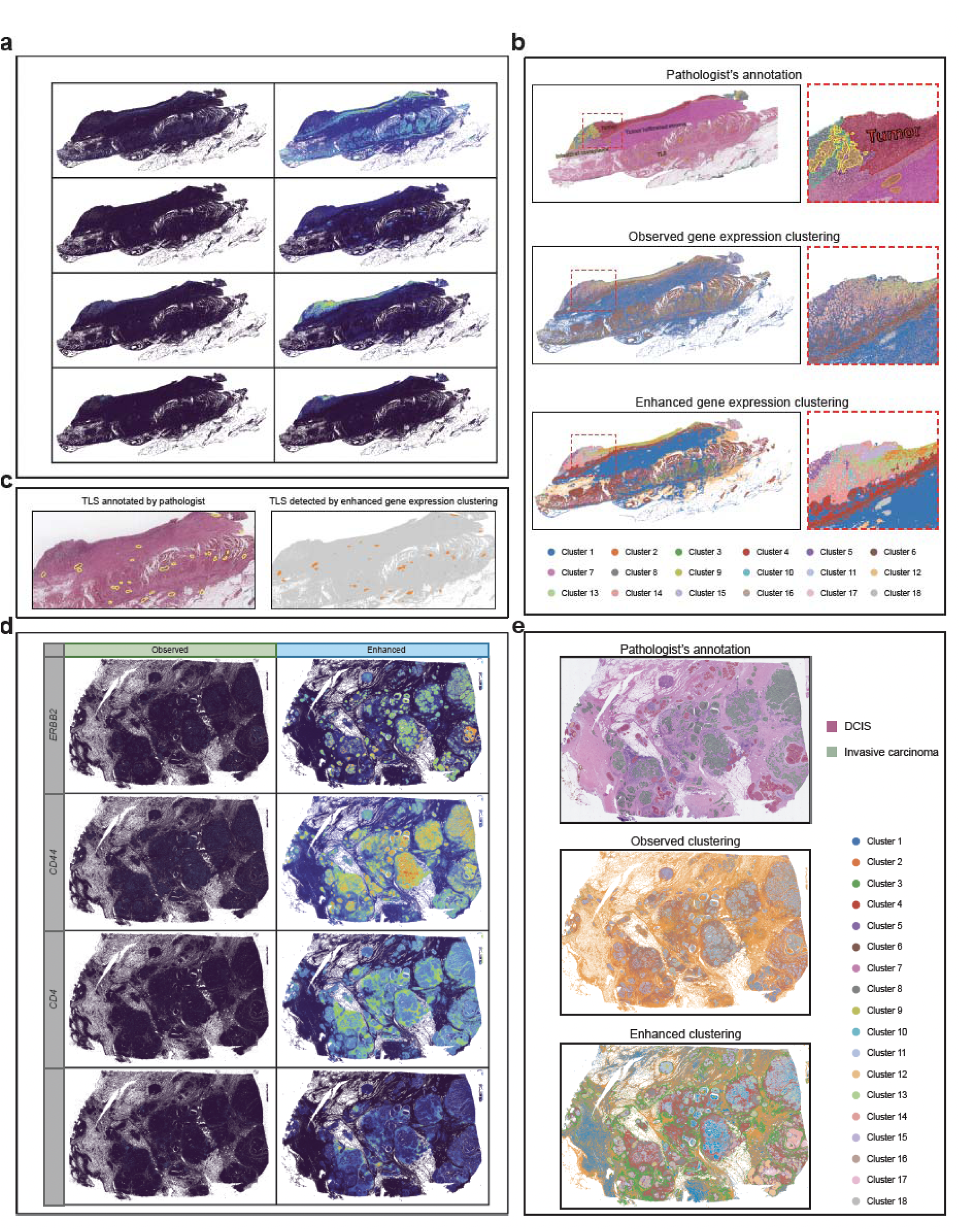
Generalization of Pixel2Gene across high-resolution spatial transcriptomics platforms and large tissue areas. **a**, Selected gene expression (*ACTA2*, *CD3D*, *EPCAM*, *KRT20*) from a Xenium gastric cancer sample. Pixel2Gene-enhanced predictions exhibit improved spatial coherence and delineate tissue boundaries and compartmentalization more clearly than the noisy observed expression. **b**, From top to bottom: pathologist annotation, clustering based on observed expression, and clustering based on Pixel2Gene-enhanced expression. In the mucosa region, observed data yields noisy and fragmented clusters, while Pixel2Gene recovers structures that align with expert annotations. **c**, Detection of tertiary lymphoid structures (TLS) in the same sample. Pathologist-annotated TLS regions (left) are successfully recovered via unsupervised clustering of the Pixel2Gene-enhanced expression (right), demonstrating the method’s ability to uncover complex immune niches. **d**, Application to a Xenium 5K breast cancer sample with noisy whole-transcriptome expression. Expression of *ERBB2*, *CD44*, *CD4*, and *MS4A1* shows clearer spatial organization after enhancement. **e**, From top to bottom: pathologist annotation, observed clustering, and clustering on Pixel2Gene-enhanced expression. Enhanced clustering resolves spatial compartments consistent with expert-labeled regions, illustrating Pixel2Gene’s generalizability to different sample types and platforms.

To further highlight Pixel2Gene’s strengths, we focused on the upper-left region of the tissue containing signet ring cells, which are known to be associated with aggressive gastric cancer, unfavorable prognosis, and pivotal for treatment decisions^27,28^. Pixel2Gene precisely delineated the interface between the poorly cohesive carcinoma area enriched for signet ring cells and the neighboring gastric mucosa, with the predicted boundary closely matching the pathologist’s manual annotation. In contrast, the observed Xenium data exhibited a noisy transition zone, lacking clearly separable clusters, which made the boundary between tissue types difficult to discern **(Fig. 4b)**. We further evaluated its capacity to identify key immune features, such as tertiary lymphoid structures (TLSs). TLSs are important markers of the tumor microenvironment, linked to stronger immune responses, enhanced anti-tumor activity, and better patient prognosis^29,30^. Notably, Pixel2Gene-based clustering successfully detected all TLSs annotated by the pathologist, underscoring its sensitivity to subtle but biologically significant immune niches (**Fig. 4c**).

We further applied Pixel2Gene to Xenium 5K, a more recent platform that expands gene coverage to ∼5,000 genes but with increased noise compared to standard Xenium with default gene panels. Using a human breast cancer sample, we analyzed marker genes including *ERBB2*, *CD44*, *CD4*, and *MS4A1* (**Fig. 4d**). While the observed expression was noisy and spatially diffuse, Pixel2Gene enhancement yielded cleaner patterns that matched known biological distributions. Correspondingly, unsupervised clustering based on enhanced data revealed sharper tumor-immune boundaries and clearer cell-type segregation, in closer agreement with pathologist annotations than clustering based on raw data (**Fig. 4e**).

Collectively, these results demonstrate Pixel2Gene’s scalability and generalizability across ST platforms with distinct resolutions, noise levels, and gene panels. From reconstructing transcriptome-wide expression in noisy, full-coverage platforms like Visium HD, or refining sparse measurements from high-resolution but limited-gene platforms like Xenium and Xenium 5K, Pixel2Gene consistently recovers biologically meaningful spatial structures. Importantly, its ability to process centimeter-scale tissue sections, such as those from gastric and breast cancer samples, illustrates its potential for whole-slide spatial analysis in both research and clinical contexts.

### Pixel2Gene reconstructs complete expression maps from data with sparse coverage

To further demonstrate the broad applicability of Pixel2Gene, we further applied it to CosMx, a high-resolution ST platform that provides subcellular resolution across multiple small, discrete targeted fields of view (FOVs). While CosMx achieves high spatial granularity, it is fundamentally constrained by its limited spatial coverage. A typical CosMx experiment profiles approximately 200 discrete FOVs (each 0.5 mm × 0.5 mm), covering only less than 5% of a standard tissue section (∼15 mm × 20 mm per slide). Expanding coverage beyond this limit is costly, time-consuming, and prone to RNA degradation and technical dropout, often resulting in drastically reduced molecule capture, with average molecule counts often dropping below 100 transcripts per cell, which compromises biological interpretability and reliable downstream analysis. More critically, even within the selected FOVs, gene expression signals are sparse, with many expressed molecules uncaptured. This severely hinders the ability to delineate tissue structures. Such limitations are especially problematic in complex tissues like the kidney, where fine-grained spatial continuity, such as that of nephron substructures including glomeruli and proximal tubules, is biologically essential but cannot be reliably inferred from fragmented gene expression measurements alone. These challenges underscore the need for computational approaches that compensate for sparse and incomplete measurements to enable the reconstruction of continuous, high-resolution gene expression maps across entire tissue sections and to improve the interpretability of CosMx datasets.

Pixel2Gene addresses these limitations by reconstructing continuous, high-resolution expression maps from sparse, noisy CosMx data. We applied the model to two kidney samples: one from a healthy donor (HK3039; **Fig. 5a-c**) and one from a patient with type 2 diabetes (HK2844; **Fig. 5d-f**). In both cases, the raw CosMx data exhibited incomplete FOV coverage, discontinuous expression patterns, and weak molecular signals. Pixel2Gene effectively filled in the large spatial gaps between FOVs while denoising the existing measurements, yielding spatially coherent gene expression landscapes across the whole tissue sections. To better illustrate this, we visualized the expression patterns of four biologically distinct marker genes: *AQP2* (collecting duct principal cells), *TAGLN* (vascular smooth muscle), *NPHS2* (podocytes in glomeruli), and *EMCN* (endothelial cells). In the original CosMx data, these markers appeared fragmented and discontinuous, making it difficult to link expression to anatomical compartments. Pixel2Gene restored their spatial organization with sharply defined regional boundaries, clearly delineating glomeruli, vascular structures, and epithelial zones with high fidelity.

**Fig. 5.**
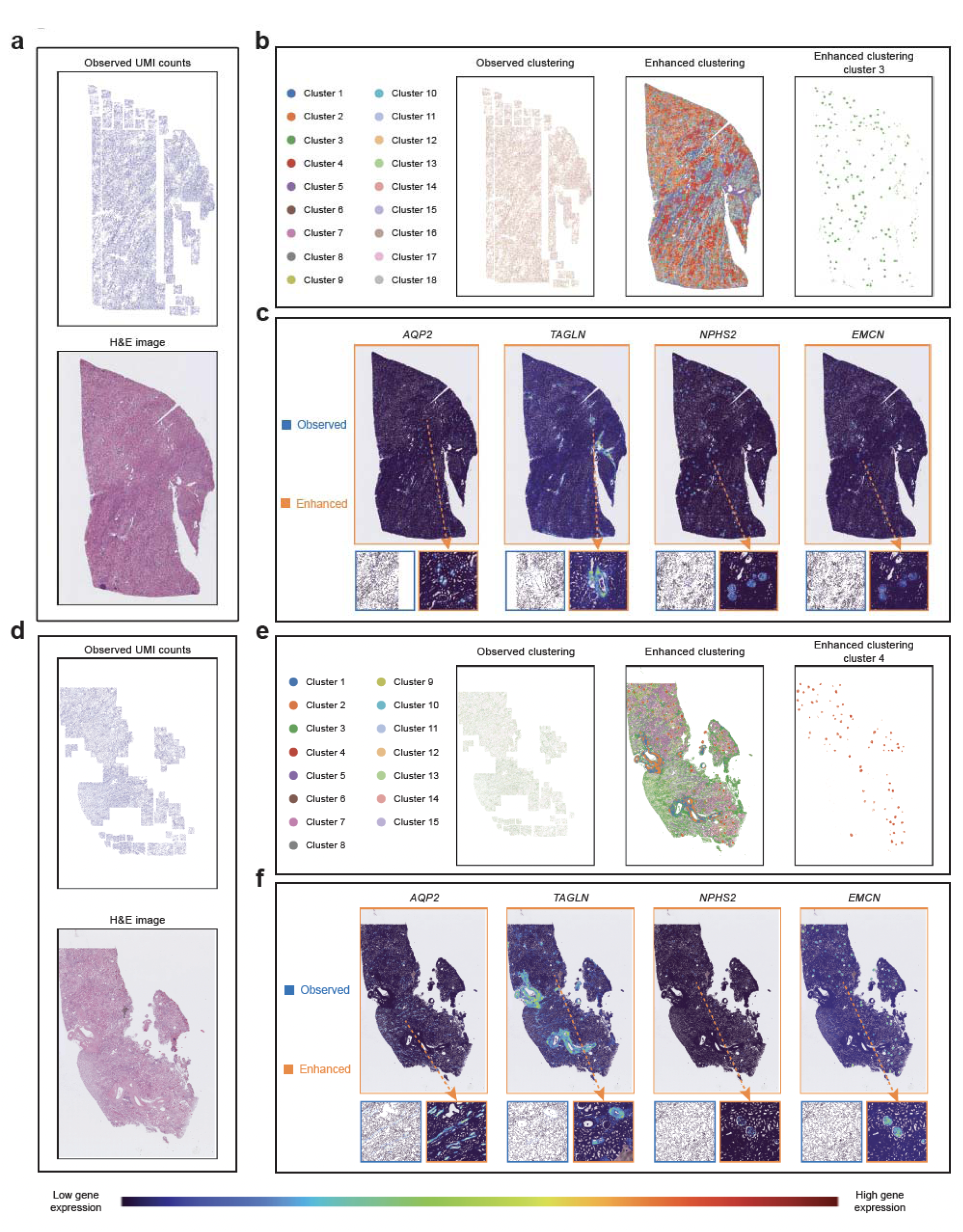
Pixel2Gene reconstructs spatial expression maps from sparse CosMx data. **a–c**, HK3039 (healthy kidney sample). **a**, Observed UMI counts across the tissue (top) and the corresponding H&E image (bottom), showing the sparse coverage typical of CosMx. **b**, Spatial clustering results from observed expression (left) and Pixel2Gene-enhanced expression (middle). Right: glomeruli identified using enhanced gene expression clustering. **c**, Expression of selected marker genes (*AQP2*, *TAGLN*, *NPHS2*, *EMCN*) in a zoomed-in region. Left: observed; right: enhanced. Pixel2Gene restores compartment-specific boundaries obscured in the raw data. **d–f**, HK2844 (type 2 diabetic kidney disease sample). **d**, Observed UMI counts (top) and H&E image (bottom). **e**, Clustering results from observed expression (left) and enhanced expression (middle). Right: glomeruli identified based on enhanced expression. **f**, Zoomed-in comparison of observed vs. enhanced expression for selected genes, showing improved spatial resolution and tissue delineation after Pixel2Gene enhancement.

In addition, unsupervised clustering based on Pixel2Gene-enhanced expression substantially outperformed clustering derived from the raw CosMx data, resolving biologically meaningful domains, such as glomeruli and proximal tubules, that were blurred or entirely missing in the original measurements. These reconstructions not only improved biological interpretability but also enabled continuous tissue segmentation and structural annotation across otherwise disconnected FOVs. Notably, Pixel2Gene’s benefits were consistent across both the healthy and diseased tissues. In the diabetic kidney sample (HK2844), where pathology-associated spatial disorganization further obscured structure in the raw CosMx data, Pixel2Gene produced interpretable expression maps that revealed preserved and altered tissue microenvironments. This highlights Pixel2Gene’s ability to bridge molecular and spatial gaps in high-resolution but low-coverage ST datasets, which empowers robust and biologically meaningful spatial analysis even in highly fragmented or degraded tissue samples.

### Pixel2Gene enables full-tissue prediction and out-of-sample prediction

Pixel2Gene not only enhances gene expression within measured regions but also supports *de novo* inference in unprofiled areas, extending spatial gene expression mapping beyond experimentally measured tissue regions. This capability is particularly useful for platforms like Visium HD, where the H&E image spans a large tissue section while gene expression is profiled in only a small 6.5 mm × 6.5 mm region. To demonstrate this, we applied Pixel2Gene to the Visium HD CRC-P2 sample, where only partial coverage was available, and reconstructed spatial gene expression across the entire tissue section using the co-registered H&E image (**Fig. 6a**). Marker genes such as *COL1A1*, a stromal collagen gene, exhibited spatially coherent expression patterns that aligned closely with histologically defined tissue compartments. Furthermore, unsupervised clustering of the full-tissue predictions revealed biologically distinct spatial domains, including the tumor core, invasive front, and normal mucosa, which were consistent with pathologist annotations. These results illustrate Pixel2Gene’s ability to extrapolate expression patterns into unmeasured tissue regions while preserving fine-grained tissue architecture and biological interpretability. Additional examples further demonstrate that Pixel2Gene consistently supports full-tissue prediction across genes (**Supplementary Fig. 4**).

**Fig. 6.**
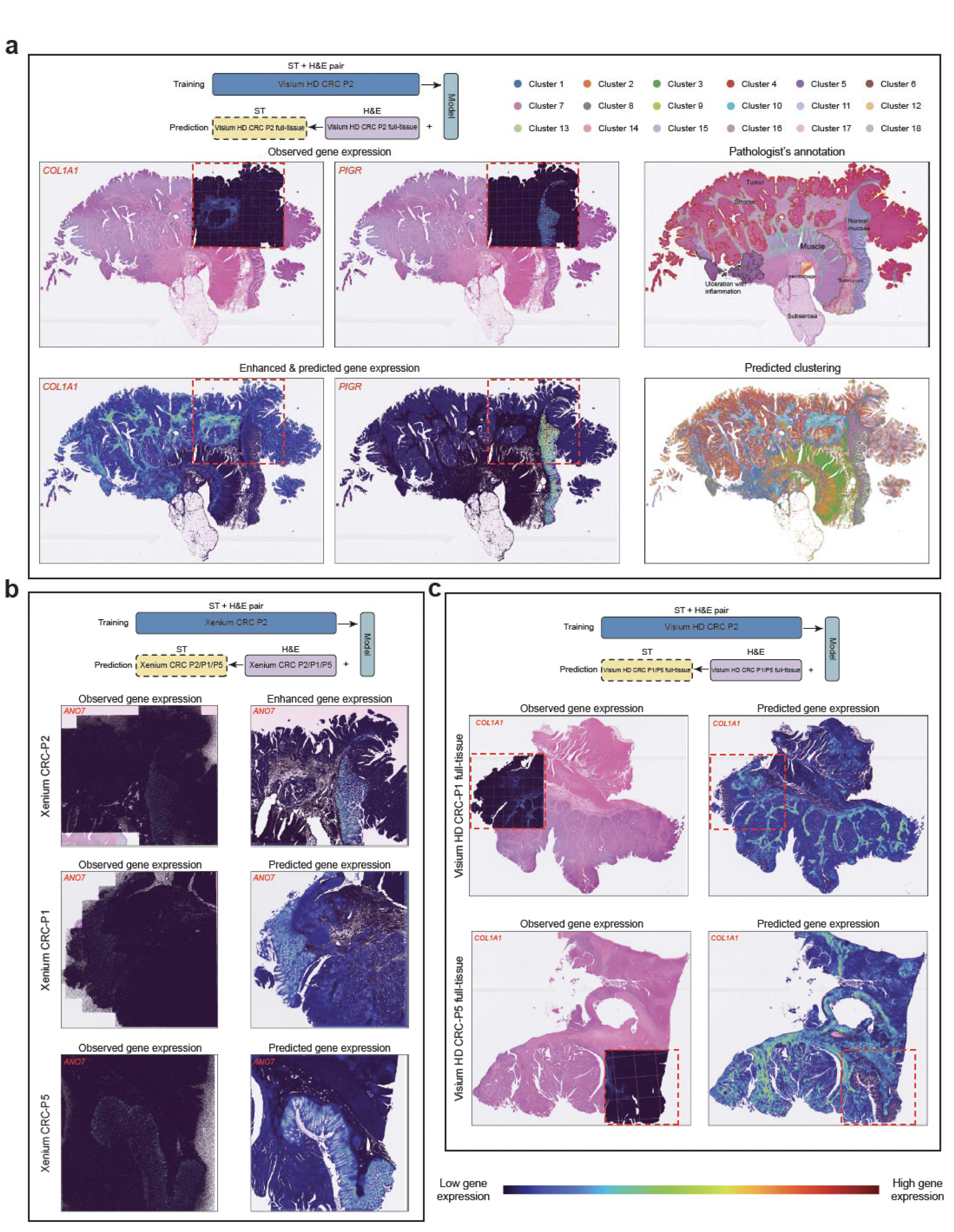
Full-tissue and out-of-sample spatial gene expression prediction using Pixel2Gene. **a**, Full-tissue prediction for Visium HD CRC-P2. Top row shows observed spatial expression of *COL1A1* (left) and *PIGR* (middle) within the profiled region, alongside the pathologist’s annotation (right). Bottom row shows Pixel2Gene-predicted full-tissue expression of *COL1A1* (left) and *PIGR* (middle) across the entire slide, together with unsupervised clustering based on the full-tissue predicted expression (right), revealing spatial domains corresponding to tumor core, invasive front, and normal mucosa. **b**, Xenium out-of-sample prediction. Pixel2Gene was trained on Xenium CRC-P2 and applied to held-out samples CRC-P1 and CRC-P5. Spatial expression of *ANO7* is shown across: Row 1 – CRC-P2 observed and Pixel2Gene-enhanced expression (training sample); Row 2 – CRC-P1 observed expression and Pixel2Gene-predicted expression; Row 3 – CRC-P5 observed expression and Pixel2Gene-predicted expression. **c**, Out-of-sample full-tissue prediction using Visium HD CRC-P2 as training. Spatial expression of *COL1A1* is shown over full-slide H&E images: top row – CRC-P1 observed (left) and Pixel2Gene-predicted (right) expression; bottom row – CRC-P5 observed (left) and predicted (right) expression.

Pixel2Gene also generalizes robustly across patients. We trained a model on Xenium data from CRC-P2 (patient 2) and applied it to two independent colorectal cancer samples, CRC-P1 (patient 1) and CRC-P5 (patient 5), profiled using the same platform but from different patients. Despite partial overlap in gene panels across the three samples, Pixel2Gene successfully predicted the full CRC-P2 gene panel in both CRC-P1 and CRC-P5, demonstrating the model’s ability to transfer learned spatial patterns across biologically similar tissues. As an illustrative example, the expression of *ANO7*, a gene present in all three panels and associated with epithelial differentiation, was consistently recovered in CRC-P1 and CRC-P5, showing clear enrichment along tumor glands that closely mirrored patterns in the training sample CRC-P2 (**Fig. 6b**). Importantly, beyond overlapping genes, Pixel2Gene also effectively inferred the expression of non-measured genes in CRC-P1 and CRC-P5 by leveraging histological features and spatial context. Consistent enhancement and out-of-sample prediction performance across additional genes is shown in **Supplementary Figs. 7-9**. These results demonstrate that Pixel2Gene not only generalizes across patients but also effectively expands the set of spatially profiled genes in each tissue beyond the limits of experimentally designed gene panels.

Finally, Pixel2Gene’s dual ability to perform out-of-sample prediction and full-tissue reconstruction unlocks substantial practical advantages. By training on a single preprocessed and well-filtered sample such as Visium HD CRC-P2, researchers can apply the model to H&E-only full-slide images from additional tissue samples (e.g., CRC-P1 and CRC-P5) without requiring further ST profiling (**Fig. 6c**). This strategy substantially reduces experimental cost and turnaround time by requiring spatial profiling of only a small, representative region, guided by experimental design tools such as S2-omics, while leveraging routine H&E images for large-scale inference, thereby eliminating the need for costly and time-consuming spatial profiling of every sample. By capitalizing on the affordability and scalability of routine H&E workflows, Pixel2Gene enables comprehensive ST analyses across large patient cohorts. This approach facilitates cost-effective and scalable spatial omics studies and opens the door to broader clinical and translational applications through systematic exploration of tissue heterogeneity at scale. Additional examples of full-tissue and out-of-sample prediction using Visium HD are provided in **Supplementary Figs. 5-6**.

### Pixel2Gene enables robust spatial pattern discovery from technically degraded post-Xenium Visium HD data

ST profiling performed after Xenium imaging provides an opportunity to obtain complementary measurements from the same tissue section but also introduces additional technical challenges. The Xenium workflow preserves overall tissue morphology and enables downstream applications, including H&E staining and post-Xenium Visium HD whole-transcriptome profiling, to be performed on the same tissue section. However, post-Xenium Visium HD data are generated after multiple rounds of in situ processing, often resulting in reduced RNA capture efficiency and increased technical noise compared with standard Visium HD experiments.

A common strategy to mitigate this limitation is the joint integration of Xenium and post-Xenium Visium HD profiles, leveraging the higher molecular resolution of Xenium to stabilize clustering and spatial pattern discovery while retaining whole-transcriptome coverage from Visium HD. However, Visium HD profiling is restricted to a selected 6.5 mm × 6.5 mm capture area, leaving substantial portions of the tissue section unmeasured. As a result, joint integration is inherently confined to regions with overlapping coverage, limiting spatial pattern discovery to Visium HD-profiled areas and preventing inference of tissue organization beyond those regions.

To assess whether Pixel2Gene can overcome the technical degradation of post-Xenium Visium HD data, we applied Pixel2Gene exclusively to post-Xenium Visium HD data from a human lung cancer sample. We then compared the resulting spatial gene expression clustering with both observed post-Xenium Visium HD clustering and clustering obtained from joint integration of Xenium and post-Xenium Visium HD data, which serves as a higher-quality reference (**Fig. 7)**. Despite using only post-Xenium Visium HD expression as input, Pixel2Gene produced spatial domains that were more coherent and spatially continuous than those derived from joint integration, with clearer alignment to histological structures observed in the corresponding H&E image.

**Fig. 7.**
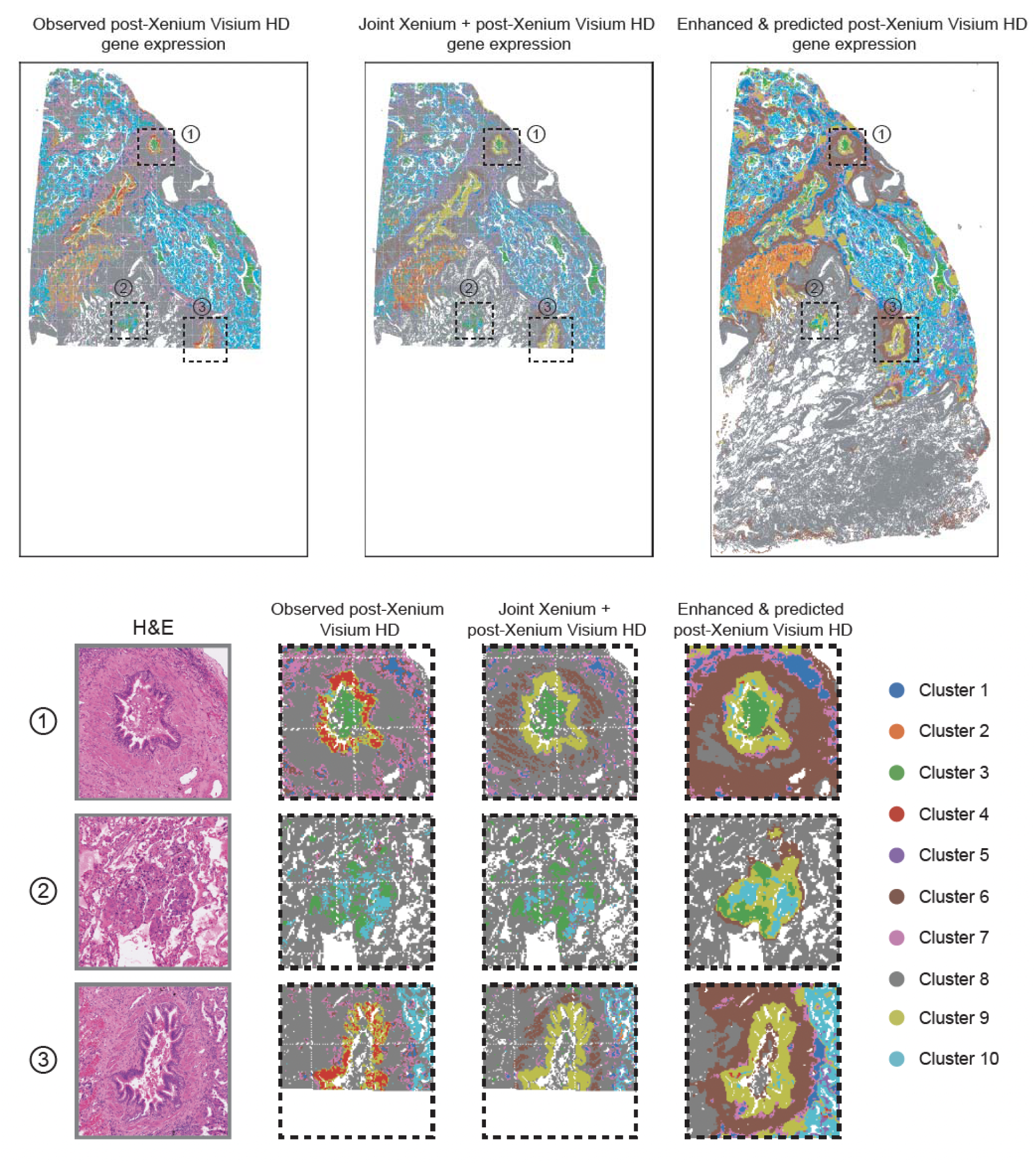
Pixel2Gene outperforms joint Xenium–Visium HD integration for post-Xenium spatial reconstruction and enables prediction in unmeasured regions. **a**, Comparison of spatial gene expression clustering across three analysis strategies applied to a post-Xenium Visium HD sample. From left to right: clustering based on observed post-Xenium Visium HD expression, clustering based on joint integration of Xenium and post-Xenium Visium HD expression, and clustering based on Pixel2Gene-enhanced and predicted post-Xenium Visium HD expression. Three regions of interest are highlighted for detailed comparison. **b–d**, Zoom-in comparisons for the three highlighted regions. For each region, panels show the corresponding H&E image, clustering from observed post-Xenium Visium HD expression, clustering from joint Xenium + post-Xenium Visium HD expression, and clustering from Pixel2Gene-enhanced and predicted post-Xenium Visium HD expression. Pixel2Gene predicts biologically plausible structures in regions lacking direct gene expression measurements, demonstrating its ability to extrapolate beyond experimentally profiled areas.

Importantly, Pixel2Gene additionally enabled prediction of spatial gene expression patterns in regions lacking direct Visium HD measurements, extending clustering and tissue compartmentalization beyond the experimentally profiled capture area. In contrast, joint integration approaches remained restricted to regions with overlapping Xenium and Visium HD coverage and could not infer structure in unmeasured tissue. These results demonstrate that Pixel2Gene can leverage histological context to denoise technically degraded post-Xenium Visium HD data and infer biologically meaningful spatial organization across entire tissue sections, achieving reconstruction quality comparable to or exceeding joint multi-platform integration while requiring only a single post-Xenium Visium HD assay.

## Discussion

Pixel2Gene addresses fundamental limitations of current high-resolution ST technologies, including data sparsity, technical noise, and incomplete gene and spatial coverage. By integrating multiscale histological features with high-quality, filtered ST measurements, Pixel2Gene reconstructs high-resolution, biologically coherent expression maps that uncover fine-grained tissue architecture often obscured in the raw data. This capability is particularly valuable in complex tissues such as tumors and kidneys, where preserving the spatial continuity of tissue structures, such as tumor invasive fronts or nephron components including glomeruli, is essential for accurate biological interpretation. Notably, these advantages persist even under technically degraded, highlighting Pixel2Gene’s robustness in challenging experimental scenarios.

A major strength of Pixel2Gene is its flexible, superpixel-based hierarchical framework. Unlike approaches that depend on explicit cell or nucleus segmentation, Pixel2Gene operates directly on H&E images, using superpixels with multiscale visual cues to offer robustness to staining variability and morphological heterogeneity. This design enables generalization across diverse ST platforms, including Visium HD, Xenium, and CosMx, and supports various tissue types and gene panel sizes without platform-specific tuning. To obtain cell-level gene expression estimates, one can calculate the weighted sum of gene expression for superpixels that overlap with a cell, an approach employed by iStar^31^.

In contrast, recent methods such as GHIST^22^, while demonstrating promising results on Xenium data, remain limited in scope and flexibility. GHIST requires nucleus segmentation as a prerequisite, introducing dependencies on nuclear staining quality and adding extra computational steps. Moreover, GHIST relies on a non-hierarchical CNN for image feature extraction, which may restrict its ability to capture multiscale tissue context. Pixel2Gene avoids these limitations by operating on a segmentation-free, superpixel-based representation, and by leveraging hierarchical feature extraction to model tissue organization across multiple spatial scales. Finally, GHIST has been demonstrated only on Xenium data, whereas Pixel2Gene generalizes robustly across multiple platforms and tissue types without reliance on cell-level annotations. These features underscore Pixel2Gene’s adaptability, scalability, and architectural advantages for comprehensive inference of spatial molecular profiles.

Beyond enhancing directly measured tissue regions, Pixel2Gene also supports out-of-sample, full-tissue prediction, where models trained on a small number of profiled regions can be applied to new samples profiled only by H&E staining. This capability substantially reduces experimental costs while expanding spatial coverage, leveraging the widespread availability and low cost of routine histology. As a result, Pixel2Gene opens new avenues for broader clinical and translational applications, including population-scale studies to prospective biomarker discovery.

Despite these strengths, Pixel2Gene’s performance depends on the quality and representativeness of its training data. Accurate inference requires training sets that capture biologically informative regions, preserve tissue morphology, and include genes with sufficient expression signal. Additionally, since ground truth expression is not available in unprofiled regions, the current performance evaluation of Pixel2Gene relies on downsampling-based benchmarks, which may not fully reflect real-world complexity. Future work may focus on optimizing training set selection, incorporating self-supervised or multimodal learning objectives, and developing an active sampling strategy for informative regions to improve generalization and robustness.

In summary, Pixel2Gene provides a robust and generalizable solution for enhancing and predicting spatial gene expression across tissue sections. Its platform-agnostic design, reliance on routine histology, and capacity for out-of-sample prediction make it well-suited for widespread adoption in biomedical research and clinical settings. By bridging histology and transcriptomics, Pixel2Gene advances scalable and interpretable spatial analyses and lays the foundation for future broader applications in precision medicine.

## Acknowledgments

M.L. was partly supported by the following NIH grants R01HG013185, R01LM014592, U19NS135582, R01HL171595, and U01CA294518.

## Author Contributions

This study was conceived of and led by M.L. S.Y. designed the model and algorithm with input from M.L., implemented the software, and led data analyses. A.S. developed the HistoSweep package for histology image quality control and high-resolution tissue masking. S.J. helped with data analysis. B.D. and K.S. provided the kidney CosMx data and advised on interpretation. S.I. annotated the breast cancer sample. J.H.P. annotated the CRC samples. T.H.H. provided the gastric cancer Xenium data. S.Y. and M.L. wrote the paper with feedback from other co-authors.

## Methods

### Data preprocessing

#### Xenium and Xenium 5K

After aligning the transcriptomic profiling data to the post-Xenium H&E pixel space, transcripts are aggregated into square spatial bins based on their transformed coordinates, with each bin corresponding to a 16 × 16-pixel region in the rescaled H&E image, referred to as a superpixel, to enable one-to-one alignment with the superpixel grid used for further analysis.

#### Rescaling

The H&E image is rescaled so that each pixel represents 0.5 µm × 0.5 µm. This standardization ensures that each 16 × 16-pixel superpixel tile corresponds to an 8 µm × 8 µm area, approximately the size of a single cell. To support hierarchical feature extraction in later stages, the rescaled H&E image is padded so that its height and width are both divisible by 256. The spatial coordinates of each gene expression bin or cell, expressed in the H&E pixel coordinate system, are rescaled using the same factor to maintain alignment between the image and the transcriptomic data. Due to technical variation, some Visium HD bins may fall partially outside the grid or overlap with others after rescaling; these bins are excluded during preprocessing to ensure consistent and reliable spatial mapping.

### Extracting hierarchical histology image features

Inspired by how pathologists examine tissues, i.e., starting with large-scale structures and zooming in to fine cellular features, we adopt a hierarchical image feature extraction framework to capture spatial patterns at multiple resolutions. This approach employs a Vision Transformer (ViT) to extract features across scales, beginning with small local patches to capture detailed morphological information, and progressively expanding to larger contextual regions to encode global tissue organization. Each 16 × 16-pixel tile ultimately results in a feature vector of length 579, containing multi-level features as well as raw RGB values. Implementation details of this multi-scale feature extraction pipeline follow the methodology described in iStar.

### Selecting high-quality spatial profiling data

To ensure only high-quality spatial profiling data are used in the training step, we apply a two-step filtering procedure that integrates both tissue morphology and gene expression quality:

#### 1. Tissue-detection masking

In this step, we aim to exclude unwanted regions such as background, debris, and low-content areas while retaining only histologically meaningful tissue. Specifically, we identify and remove (i) background and acellular regions lacking sufficient stain or structural content, and (ii) visually irregular, potentially damaged, or noisy regions that may still contain some stain but are unlikely to yield reliable data. This is accomplished by analyzing both stain density and texture features across the tissue, resulting in a binary mask that preserves dense, textured, and well-stained tissue regions.

#### 2. Spatial profiling-quality masking

Beyond histological relevance, we also prioritize regions with robust gene expression capture. To achieve this, we first perform K-means clustering on superpixel-level histological features to account for tissue heterogeneity. Within each cluster, we filter out superpixels whose gene expression profiles, as represented by total UMI counts, fall below a specified quantile threshold (e.g., bottom 30%), thereby removing regions likely to contain low-quality or noisy molecular data.

By combining these two masks, we ensure that downstream analysis is performed only on regions that are both histologically valid and molecularly informative, thereby improving the reliability and interpretability of spatial transcriptomic results.

### Predicting super-resolution gene expression in large tissues

The previously developed iStar method enhances spatial resolution for Visium data by utilizing a single ST sample as both input and training data. In contrast, our setting involves ST samples from various platforms that already provide single-cell resolution, allowing for a one-to-one correspondence between each cell bin and its associated image superpixel.

To leverage high-quality ST data, we incorporate ST samples with reliable gene expression profiling to perform both in-sample super-resolution enhancement and out-of-sample full-tissue prediction. Given an input histology image, histological features are extracted as

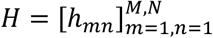

where ℎ*_mn_* represents the feature at superpixel (*m*, *n*). Let *X_kmn_* denote the observed expression of gene *k* at superpixel (*m*, *n*) in the training ST sample, and let *P* be the set of superpixels with measured gene expression. Then for each gene *k*, a prediction model *g_k_*(·) is trained by minimizing the mean squared error (MSE) loss between predicted and observed gene expression values:

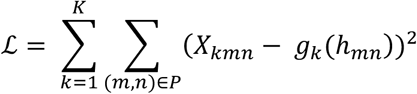

After training, gene expression at each superpixel (*m*, *n*) ∈ *P* of the training sample is predicted as

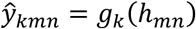

effectively leveraging information from all training superpixels across the entire tissue image, yielding enhanced super-resolution gene expression within the profiled region.

The trained model can further be applied to all superpixels across the entire tissue image, yielding a continuous gene expression prediction

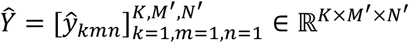

including regions without measured expression. Moreover, the same model can be directly applied to histologically similar regions in new, unseen tissue samples to perform out-of-sample expression prediction.

### Neural network architecture and hyperparameter tuning

We employed a multi-layer perceptron (MLP) architecture consisting of a feed-forward neural network with four hidden layers, each containing 256 neurons. A leaky ReLU activation function with a negative slope of α = 0.1 was applied to each hidden layer to allow for non-linearity while avoiding neuron inactivity. The model was trained using the Adam optimizer with a batch size of 128 for 1,200 epochs. The learning rate was adjusted based on the size of the input training dataset to ensure stable convergence across samples of varying coverage.

### Downsampling simulation

Due to the inherent noise and incompleteness of single-cell resolution ST data generated from Visium HD, ground truth expression values are not directly available. Therefore, we restricted the evaluation to the top 300 highly expressed genes in the Visium HD platform, as these genes are less affected by dropout events and provide a more stable reference for assessing reconstruction accuracy.

The downsampling procedure was implemented as follows:

#### 1. Shrinkage factor generation

For each superpixel *i*, we independently sample a shrinkage factor *τ_i_* from a shared Gamma distribution, enabling systematic yet heterogeneous coverage reduction across the spatial domain,

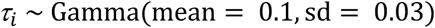

#### 2. Library size adjustment

The original total UMI count for each superpixel *i*, *N_i_*, is multiplied by its corresponding shrinkage factor to generate a new, reduced library size:

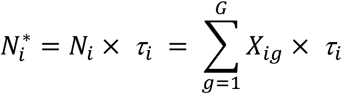

#### 3. Library size adjustment

For each superpixel, we generate synthetic counts by sampling from a multinomial distribution:

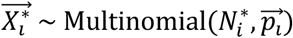

where the probability vector 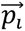 represents the relative abundance of each gene in the original gene expression count data.

### Evaluation metrics

For the downsampling experiments, we assessed reconstruction accuracy by comparing the predicted gene expression profiles to the original (pre-downsampled) ground truth across spatial locations. To capture different aspects of agreement between the downsampled, enhanced, and ground truth data, we compute a comprehensive set of evaluation metrics. These included Pearson and Spearman correlation coefficients, which quantify linear and rank-based concordance, respectively; cosine similarity, which assesses directional alignment between normalized expression vectors; and error-based metrics such as root mean squared error (RMSE) and mean absolute error (MAE). All expression values are normalized to the [0, 1] range prior to evaluation, and metrics are computed either across all spatial spots or restricted to regions with nonzero ground truth expression.

To evaluate spatial autocorrelation at multiple resolutions, we evaluate spatial autocorrelation at multiple resolutions using Moran’s *I*, a statistic that measures how spatially clustered or dispersed the expression signal is, with a fast convolution-based method that handles missing values. To simulate different spatial scales, we applied sum pooling to the expression matrix using window sizes corresponding to 1× (no pooling), 2×, and 4× pooling factors. These correspond to neighborhood radii of 16 pixels, 32 pixels, and 64 pixels, respectively, and are reported as Moran’s *I*-16, Moran’s *I*-32, and Moran’s *I*-64. At each scale, the pooled expression matrix is passed to the Moran’s *I* function to measure the degree of local spatial autocorrelation. This multi-scale approach captures both fine-grained and broader spatial continuity, offering a robust view of how well the reconstructed data preserves biologically relevant spatial structure.

### Spatial clustering analysis

We perform spatial clustering in a two-stage manner to ensure both stability and interpretability of the resulting clusters. First, we apply MiniBatch K-means clustering to the superpixel-level data, intentionally specifying a slightly higher number of clusters than the desired number. This over-partitioning ensures that fine-grained spatial structures are captured early in the process.

Next, we refine the clustering results using an iterative hierarchical clustering-based merging strategy. Specifically, we construct a hierarchical tree based on the centroids of the initial K-means clusters using a linkage matrix to merge clusters based on a dynamically adjusted distance threshold. We then further merge small clusters whose superpixel counts fall below a predefined fraction of the total counts (e.g., 0.5%). Samples from these small clusters are reassigned to their nearest larger clusters based on Euclidean distances in feature space.

We repeat this process, adjusting the threshold at which the hierarchical clustering dendrogram is cut to produce flat clusters, until the number of resulting clusters closely matches the target number. In this sense, the hierarchical merging serves as a flexible post-processing step that balances between over-segmentation and under-segmentation, producing spatial clusters that are both meaningful and size regularized.

## Data Availability

We analyzed the following datasets in this study: (1) 10x Genomics human colorectal cancer Visium HD data, P1 CRC, P2 CRC, P5 CRC (https://www.10xgenomics.com/products/visium-hd-spatial-gene-expression/dataset-human-crc). (2) the corresponding 10x Genomics Xenium human colorectal cancer datasets for CRC-P1, CRC-P2, and CRC-P5, available from the same data source. (3) 10x Genomics mouse brain Visium HD data, fresh frozen (https://www.10xgenomics.com/datasets/visium-hd-cytassist-gene-expression-mouse-brain-fresh-frozen). (4) 10x Genomics human gastric cancer tumor sample Xenium data (Gastric Patient 1, Tumor, IRB No. 30-2023-1) generated by Tae Hyun Hwang lab (DOI: 10.5281/zenodo.15164980). (5) 10x Genomics human breast cancer Xenium Prime 5K data (https://www.10xgenomics.com/datasets/xenium-prime-ffpe-human-breast-cancer). (6) human kidney CosMx data generated by Katalin Susztak (https://zenodo.org/records/17228449). (7) 10x Genomics human lung cancer Xenium data (https://www.10xgenomics.com/datasets/xenium-human-lung-cancer-post-xenium-technote) and post-Xenium Visium HD data(https://www.10xgenomics.com/datasets/visium-hd-cytassist-gene-expression-human-lung-cancer-post-xenium-expt). Details of the datasets analyzed in this paper are described in **Supplementary Table 1**.

## Code Availability

Pixel2Gene was implemented in Python and is available on GitHub at https://github.com/yaosicong1999/Pixel2Gene.

## References

1 Marx, V. Method of the Year: spatially resolved transcriptomics. Nat Methods 18, 9–14 (2021). 10.1038/s41592-020-01033-y

2 Holt, C. E. & Bullock, S. L. Subcellular mRNA Localization in Animal Cells and Why It Matters. Science 326, 1212–1216 (2009). 10.1126/science.1176488

3 Arora, R. et al. Spatial transcriptomics reveals distinct and conserved tumor core and edge architectures that predict survival and targeted therapy response. Nat Commun 14, 5029 (2023). 10.1038/s41467-023-40271-4

4 He, S. et al. High-plex imaging of RNA and proteins at subcellular resolution in fixed tissue by spatial molecular imaging. Nat Biotechnol 40, 1794–1806 (2022). 10.1038/s41587-022-01483-z

5 Moffitt, J. R. et al. Molecular, spatial, and functional single-cell profiling of the hypothalamic preoptic region. Science 362, eaau5324 (2018). 10.1126/science.aau5324

6 Oliveira, M. F. D. et al. High-definition spatial transcriptomic profiling of immune cell populations in colorectal cancer. Nat Genet 57, 1512–1523 (2025). 10.1038/s41588-025-02193-3

7 Janesick, A. et al. High resolution mapping of the tumor microenvironment using integrated single-cell, spatial and in situ analysis. Nat Commun 14, 8353 (2023). 10.1038/s41467-023-43458-x

8 Zhang, L. et al. Clinical and translational values of spatial transcriptomics. Sig Transduct Target Ther 7, 111 (2022). 10.1038/s41392-022-00960-w

9 Lim, H. J., Wang, Y., Buzdin, A. & Li, X. A practical guide for choosing an optimal spatial transcriptomics technology from seven major commercially available options. BMC Genomics 26, 47 (2025). 10.1186/s12864-025-11235-3

10 Ghaznavi, F., Evans, A., Madabhushi, A. & Feldman, M. Digital Imaging in Pathology: Whole-Slide Imaging and Beyond. Annu. Rev. Pathol. Mech. Dis. 8, 331–359 (2013). 10.1146/annurev-pathol-011811-120902

11 MD Anderson Cancer Center. Service Price List. (Research Histology Core Laboratory, MD Anderson Cancer Center, 2019). https://www.mdanderson.org/content/dam/mdanderson/documents/core-facilities/Research%20Histology%20Core%20Laboratory%20(RHCL)/RHCL%20Service%20Prices%20Fina%20III%2005.20.19.pdf

12 Roche Canada. H&E and Special Stains. https://www.rochecanada.com/solutions/diagnostics-solutions/laboratories/pathology-lab/he-and-special-stains (2024).

13 Schmauch, B. et al. A deep learning model to predict RNA-Seq expression of tumours from whole slide images. Nat Commun 11, 3877 (2020). 10.1038/s41467-020-17678-4

14 He, B. et al. Integrating spatial gene expression and breast tumour morphology via deep learning. Nat Biomed Eng 4, 827–834 (2020). 10.1038/s41551-020-0578-x

15 Bergenstråhle, L. et al. Super-resolved spatial transcriptomics by deep data fusion. Nat Biotechnol 40, 476–479 (2022). 10.1038/s41587-021-01075-3

16 Jia, Y., Liu, J., Chen, L., Zhao, T. & Wang, Y. THItoGene: a deep learning method for predicting spatial transcriptomics from histological images. Briefings in Bioinformatics 25, bbad464 (2023). 10.1093/bib/bbad464

17 Lin, Y. et al. A contrastive learning approach to integrate spatial transcriptomics and histological images. Computational and Structural Biotechnology Journal 23, 1786–1795 (2024). 10.1016/j.csbj.2024.04.039

18 Coleman, K. et al. Resolving tissue complexity by multimodal spatial omics modeling with MISO. Nat Methods 22, 530–538 (2025). 10.1038/s41592-024-02574-2

19 Coleman, K., Schroeder, A. & Li, M. Unlocking the power of spatial omics with AI. Nat Methods 21, 1378–1381 (2024). 10.1038/s41592-024-02363-x

20 Schroeder, A. et al. Scaling up spatial transcriptomics for large-sized tissues: uncovering cellular-level tissue architecture beyond conventional platforms with iSCALE. Nat Methods 22, 1911–1922 (2025). 10.1038/s41592-025-02770-8

21 Yuan, M. et al. Smart spatial omics (S2-omics) optimizes region of interest selection to capture molecular heterogeneity in diverse tissues. Nat Cell Biol 27, 2225–2238 (2025). 10.1038/s41556-025-01811-w

22 Fu, X. et al. Spatial gene expression at single-cell resolution from histology using deep learning with GHIST. Nat Methods 22, 1900–1910 (2025). 10.1038/s41592-025-02795-z

23 Wan, X. et al. Integrating spatial and single-cell transcriptomics data using deep generative models with SpatialScope. Nat Commun 14, 7848 (2023). 10.1038/s41467-023-43629-w

24 Zhao, C. et al. Innovative super-resolution in spatial transcriptomics: a transformer model exploiting histology images and spatial gene expression. Briefings in Bioinformatics 25, bbae052 (2024). 10.1093/bib/bbae052

25 Schroeder, A. et al. HistoSweep enables cellular-resolution tissue quality control for gigapixel images in digital pathology and spatial omics. bioRxiv, 2026.2001.2030.702675 (2026). 10.64898/2026.01.30.702675

26 Hwang Lab. Gastric cancer Xenium sample – BS06-9313-8_Tumor. Zenodo 10.5281/zenodo.15164980 (2025).

27 Pernot, S. Signet-ring cell carcinoma of the stomach: Impact on prognosis and specific therapeutic challenge. WJG 21, 11428 (2015). 10.3748/wjg.v21.i40.11428

28 Alcindor, T. et al. Phase 2 trial of perioperative chemo-immunotherapy for gastro-esophageal adenocarcinoma: The role of M2 macrophage landscape in predicting response. Cell Reports Medicine 6, 102045 (2025). 10.1016/j.xcrm.2025.102045

29 Weng, Y. et al. The impact of tertiary lymphoid structures on tumor prognosis and the immune microenvironment in non-small cell lung cancer. Sci Rep 14, 16246 (2024). 10.1038/s41598-024-64980-y

30 Peyraud, F. et al. Tertiary lymphoid structures and cancer immunotherapy: From bench to bedside. Med 6, 100546 (2025). 10.1016/j.medj.2024.10.023

31 Zhang, D. et al. Inferring super-resolution tissue architecture by integrating spatial transcriptomics with histology. Nat Biotechnol 42, 1372–1377 (2024). 10.1038/s41587-023-02019-9

