## Supplementary material for "Pixel2Gene enables histology-guided reconstruction and prediction of spatial gene expression"

**Supplementary Table 1: Datasets analyzed in this paper.** Spatial units correspond to image-aligned superpixels prior to any filtering or aggregation.

| Species | Tissue | Protocol | Dimensions of Dataset | Data Source |
|---|---|---|---|---|
| Human | Colorectal cancer P2 (CRC-P2) | 10x Visium HD | 545571 spatial units<br>18085 genes | https://www.10xgenomics.com/platforms/visium/product-family/dataset-human-crc |
| Human | Colorectal cancer P2 (CRC-P2) | 10x Xenium | 634586 spatial units<br>422 genes | https://www.10xgenomics.com/platforms/visium/product-family/dataset-human-crc |
| Human | Gastric cancer | 10x Xenium | 1971688 spatial units<br>377 genes | https://zenodo.org/records/15164980 |
| Human | Breast cancer | 10x Xenium 5K | 1562526 spatial units<br>5122 genes | https://www.10xgenomics.com/datasets/xenium-prime-ffpe-human-breast-cancer |
| Human | Kidney cancer HK2844 | NanoString CosMx | 195593 spatial units<br>1000 genes | https://zenodo.org/records/17228449 |
| Human | Kidney cancer HK3039 | NanoString CosMx | 205700 spatial units<br>1000 genes | https://zenodo.org/records/17228449 |
| Human | Colorectal cancer P1 (CRC-P1) | 10x Visium HD | 507680 spatial units<br>18085 genes | https://www.10xgenomics.com/platforms/visium/product-family/dataset-human-crc |
| Human | Colorectal cancer P5 (CRC-P5) | 10x Visium HD | 541958 spatial units<br>18085 genes | https://www.10xgenomics.com/platforms/visium/product-family/dataset-human-crc |
| Human | Colorectal cancer P1 (CRC-P1) | 10x Xenium | 574176 spatial units<br>422 genes | https://www.10xgenomics.com/platforms/visium/product-family/dataset-human-crc |
| Human | Colorectal cancer P5 (CRC-P5) | 10x Xenium | 635286 spatial units<br>422 genes | https://www.10xgenomics.com/platforms/visium/product-family/dataset-human-crc |
| Human | Lung cancer | 10x Xenium | 813889 spatial units<br>422 genes | https://www.10xgenomics.com/datasets/xenium-human-lung-cancer-post-xenium-technote |
| Human | Lung cancer (post-Xenium) | 10x Visium HD | 443637 spatial units<br>18049 genes | https://www.10xgenomics.com/datasets/visium-hd-cytassist-gene-expression-human-lung-cancer-post-xenium-expt |
| Mouse | Brain (Fresh Frozen) | 10x Visium HD | 453459 spatial units<br>19059 genes | https://www.10xgenomics.com/datasets/visium-hd-cytassist-gene-expression-mouse-brain-fresh-frozen |

**Supplementary Table 2: Computational cost of datasets analyzed in this paper.**
All experiments were run on NVIDIA A100 GPUs (80 GB memory) with a fixed training schedule of 600 epochs for reporting purposes. CPU usage is reported as CPU core-minutes (sum of user and system CPU time) to account for the variable number of CPU cores allocated by the scheduler. For 10-fold cross-validation, training and prediction costs are reported as the average per fold. Peak memory usage corresponds to the maximum resident set size (RSS) reported by the job scheduler, and peak GPU memory is the maximum GPU memory allocated during execution. The preprocessing stage includes image rescaling, tissue masking, and feature extraction. Preprocessing and combining steps were executed once per dataset. These measurements are provided for illustrative purposes only and are not intended as formal performance benchmarks. Runtime and resource usage may vary depending on hardware configuration, software versions, scheduler policies, I/O performance, and dataset-specific characteristics.

| dataset | step | CPU cost (core-mins) | Peak RAM (GB) | Peak GPU memory (GB) |
|---|---|---|---|---|
| Visium HD CRC-P2 | preprocess | 262.8 | 69.55 | 20.69 |
| | 10-fold training (per-fold average) | 174.07 | 83.35 | 17.33 |
| | 10-fold prediction (per-fold average) | 4.23 | 4.98 | 23.08 |
| | combining | 50.97 | 0.15 | 0.00 |
| Visium HD CRC-P1 (prediction only) | preprocess | 204.85 | 43.57 | 20.22 |
| | 10-fold prediction (per-fold average) | 7.19 | 5.15 | 51.41 |
| | combining | 53.90 | 0.15 | 0.00 |
| Visium HD CRC-P5 (prediction only) | preprocess | 206.98 | 63.39 | 21.43 |
| | 10-fold prediction (per-fold average) | 9.28 | 5.25 | 19.07 |
| | combining | 55.18 | 0.14 | 0.00 |
| Visium HD Human Lung Cancer | preprocess | 323.28 | 27.58 | 25.56 |
| | 10-fold training (per-fold average) | 194.34 | 14.31 | 36.84 |
| | 10-fold prediction (per-fold average) | 2.20 | 6.85 | 20.07 |
| | combining | 16.45 | 8.96 | 0.00 |
| Visium HD | preprocess | 198.58 | 67.14 | 41.67 |

| | | | | |
|---|---|---|---|---|
| CRC-P2 Downsampled | 10-fold training (per-fold average) | 239.25 | 81.93 | 41.10 |
| | 10-fold prediction (per-fold average) | 5.07 | 4.88 | 50.63 |
| | combining | 38.65 | 0.14 | 0.00 |
| Visium HD Mouse Brain Downsampled | preprocess | 183.14 | 66.11 | 48.27 |
| | 10-fold training (per-fold average) | 226.62 | 109.4 | 64.38 |
| | 10-fold prediction (per-fold average) | 5.20 | 4.54 | 50.58 |
| | combining | 65.36 | 0.12 | 0.00 |
| Xenium Human Gastric Cancer | preprocess | 556.2 | 31.57 | 41.67 |
| | 10-fold training (per-fold average) | 418.39 | 35.00 | 65.60 |
| | 10-fold prediction (per-fold average) | 2.64 | 24.67 | 54.03 |
| | combining | 6.64 | 0.33 | 0.00 |
| Xenium 5K Human Breast Cancer | preprocess | 551.32 | 96.71 | 55.07 |
| | 10-fold training (per-fold average) | 503.88 | 122.43 | 55.11 |
| | 10-fold prediction (per-fold average) | 6.48 | 17.05 | 50.49 |
| | combining | 30.59 | 0.26 | 0.00 |
| Xenium CRC-P2 | preprocess | 180.52 | 16.37 | 20.41 |
| | 10-fold training (per-fold average) | 120.45 | 8.00 | 23.95 |
| | 10-fold prediction (per-fold average) | 1.02 | 5.42 | 17.49 |
| | combining | 3.38 | 0.14 | 0.00 |
| Xenium CRC-P1 (prediction only) | preprocess | 188.27 | 15.79 | 21.42 |
| | 10-fold prediction (per-fold average) | 1.00 | 5.25 | 18.64 |
| | combining | 3.33 | 0.14 | 0.00 |
| Xenium CRC-P5 | preprocess | 199.10 | 15.07 | 21.43 |

| | | | | |
|---|---|---|---|---|
| (prediction only) | 10-fold prediction (per-fold average) | 0.93 | 5.09 | 17.10 |
| | combining | 3.27 | 0.13 | 0.00 |
| CosMx HK2844 | preprocess | 711.22 | 38.00 | 30.12 |
| | 10-fold training (per-fold average) | 138.4 | 36.95 | 50.44 |
| | 10-fold prediction (per-fold average) | 4.93 | 29.7 | 42.36 |
| | combining | 15.5 | 0.38 | 0.00 |
| CosMx HK3039 | preprocess | 554.62 | 26.77 | 41.01 |
| | 10-fold training (per-fold average) | 143.29 | 26.84 | 68.65 |
| | 10-fold prediction (per-fold average) | 3.35 | 20.98 | 42.12 |
| | combining | 10.59 | 0.31 | 0.00 |

**Supplementary Fig. 1:** Evaluation of Pixel2Gene enhancement in the simulated Mouse Brain data. **a,** Visualization of spatial expression patterns of selected genes in simulated mouse brain Visium HD data at 8 μm resolution, shown for the observed, downsampled, and Pixel2Gene-enhanced (downsampled-enhanced) gene expression. **b,** Quantitative comparison of prediction accuracy and spatial autocorrelation across the top 300 highly expressed genes, using the same evaluation metrics as in **Fig. 2**. **c,** Comparison of spatial clustering results across different input modalities. From left to right: Allen Brain Atlas anatomical annotation[1] (serving as a reference), clustering based on observed gene expression, downsampled gene expression, and Pixel2Gene-enhanced expression. Cluster labels were aligned to the observed clustering using a one-to-one assignment, and NMI scores with respect to the observed clustering are shown.

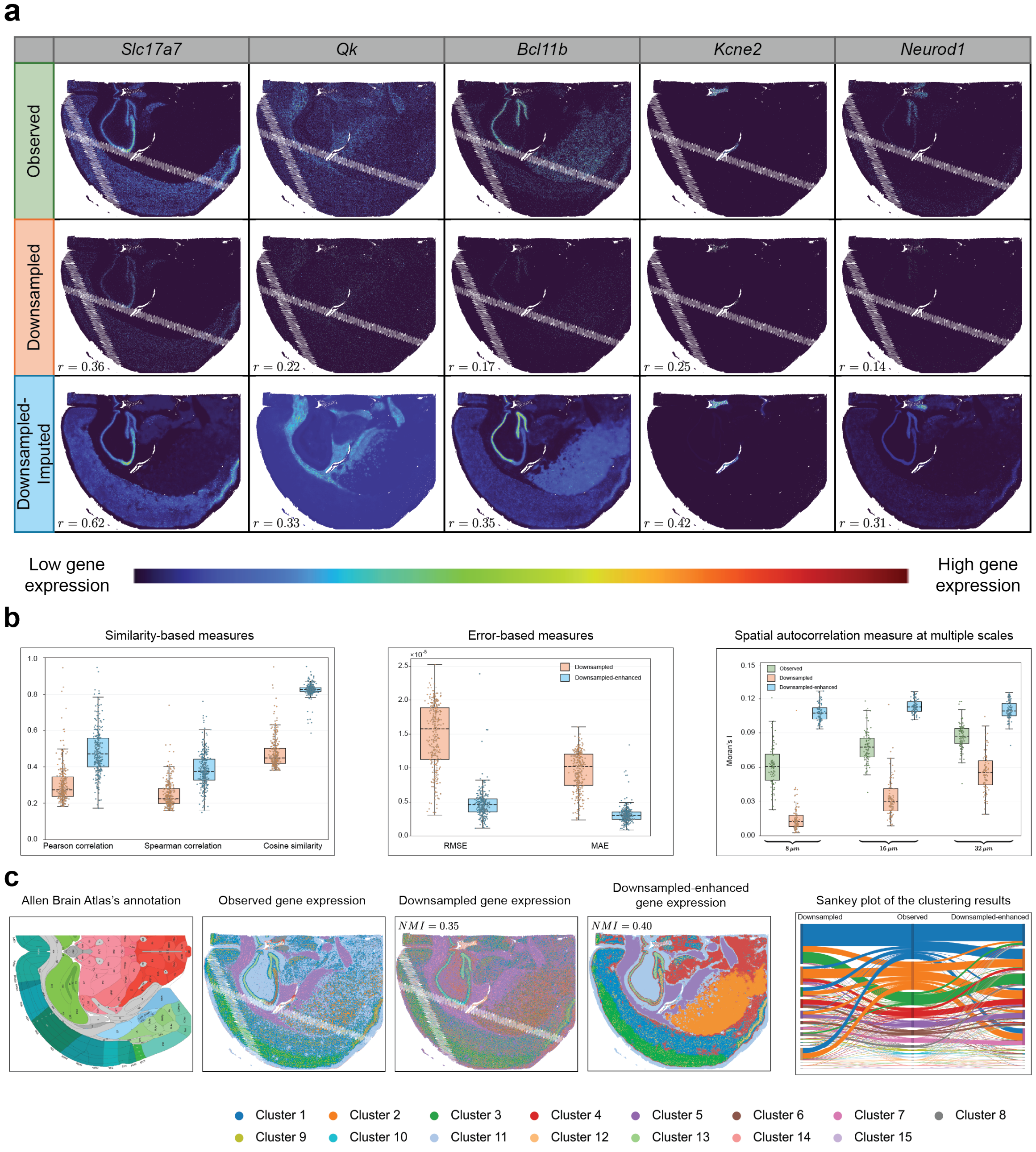

**Supplementary Fig. 2:** Pixel2Gene improves intra-cluster correlations in downsampling simulations. **a**, Boxplots of ICC distributions computed across clusters and the top 50 PCs derived from the top 300 highly expressed genes used in the CRC-P2 downsampling simulation. ICCs quantify the ratio of between-cluster to total variance per PC. Pixel2Gene-enhanced expression exhibits higher ICC values than both observed and downsampled data (mean ICC = 0.2549 versus 0.1309 and 0.1368, respectively), indicating improved intracluster coherence. **b,** Same analysis applied to simulated mouse brain Visium HD data. Pixel2Gene-enhanced expression similarly achieves higher ICC values than observed and downsampled data (mean ICC = 0.3658 versus 0.2158 and 0.1312, respectively).

**a**

ICC distributions across all clusters
in the CRC-P2 downsampling simulation

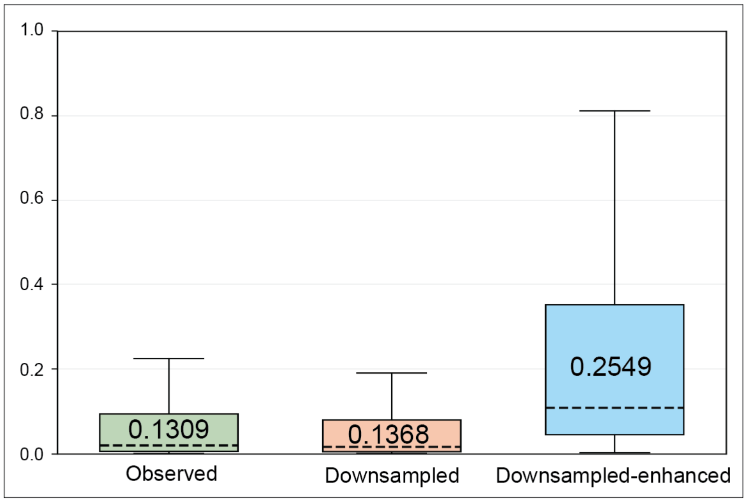


**b**

ICC distributions across all clusters
in the mouse brain downsampling simulation

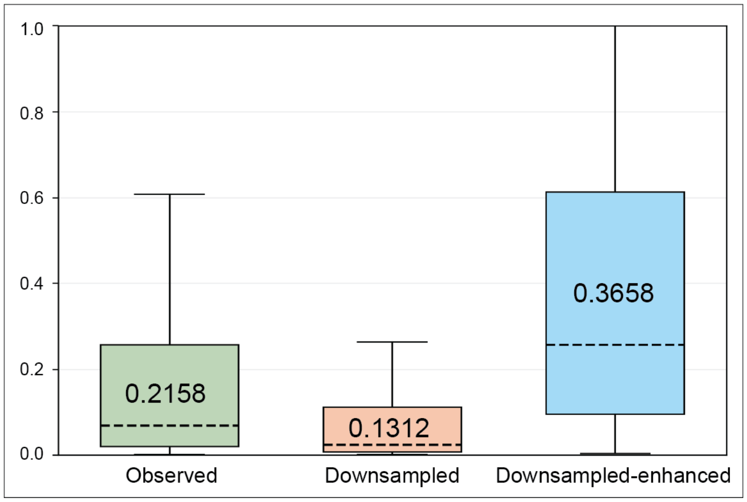

**Supplementary Fig. 3:** Additional examples of Pixel2Gene-enhanced spatial expression patterns in Visium HD CRC-P2 data. Visualization of five representative genes (*AXIN2*, *C3*, *CLCA1*, *EGFR*, and *ETS1*) from CRC-P2 Visium HD data that were not included among the top 300 highly expressed genes. For each gene, expression patterns are shown for the observed Visium HD data (left), Pixel2Gene-enhanced predictions (middle), and matched adjacent Xenium measurements (right). Pixel2Gene enhancement produces sharper and more spatially coherent expression patterns that more closely resemble high-resolution Xenium references, despite substantial noise in the original Visium HD measurements.

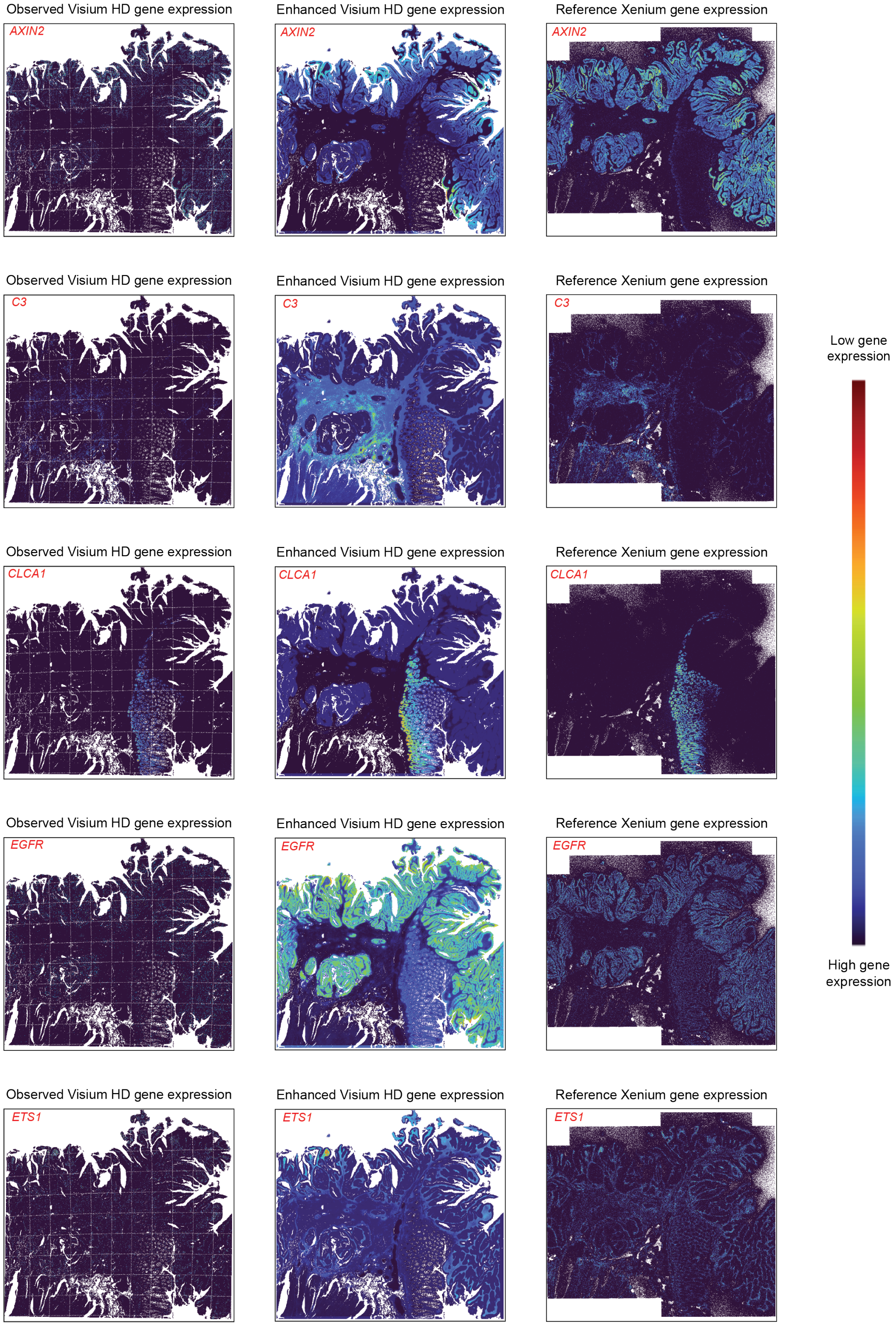

**Supplementary Fig. 4:** Additional examples of full-tissue prediction using Visium HD CRC-P2. Pixel2Gene full-tissue prediction for five additional genes in Visium HD CRC-P2. For each gene, observed expression within the profiled region is shown alongside Pixel2Gene-predicted expression across the full tissue.

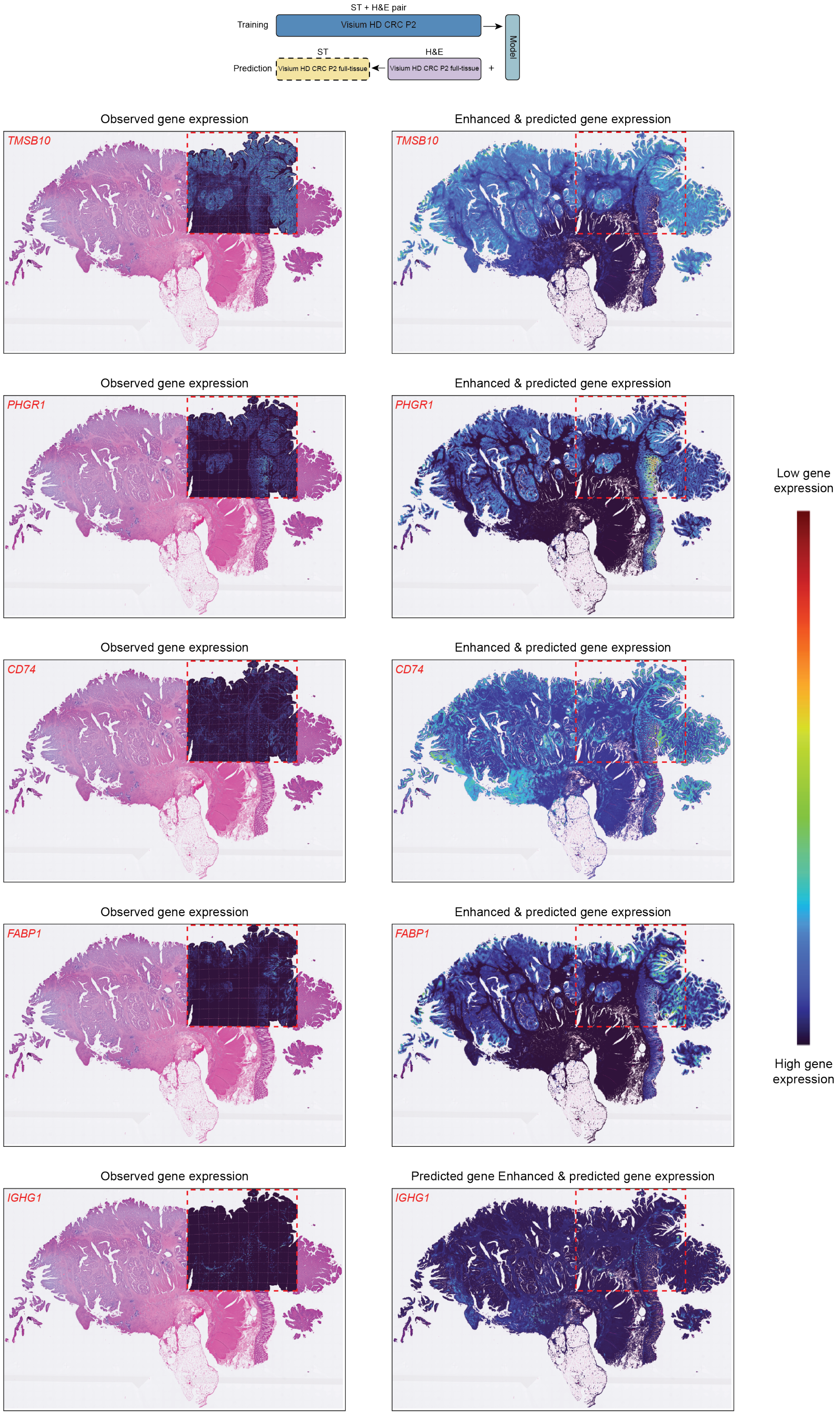

**Supplementary Fig. 5:** Additional examples of out-of-sample full-tissue prediction in CRC-P1. Pixel2Gene was trained on Visium HD CRC-P2 and applied to Visium HD CRC-P1. Full-tissue predicted expression for the same five genes shown in **Supplementary Fig. 4** is displayed alongside observed expression within the profiled region.

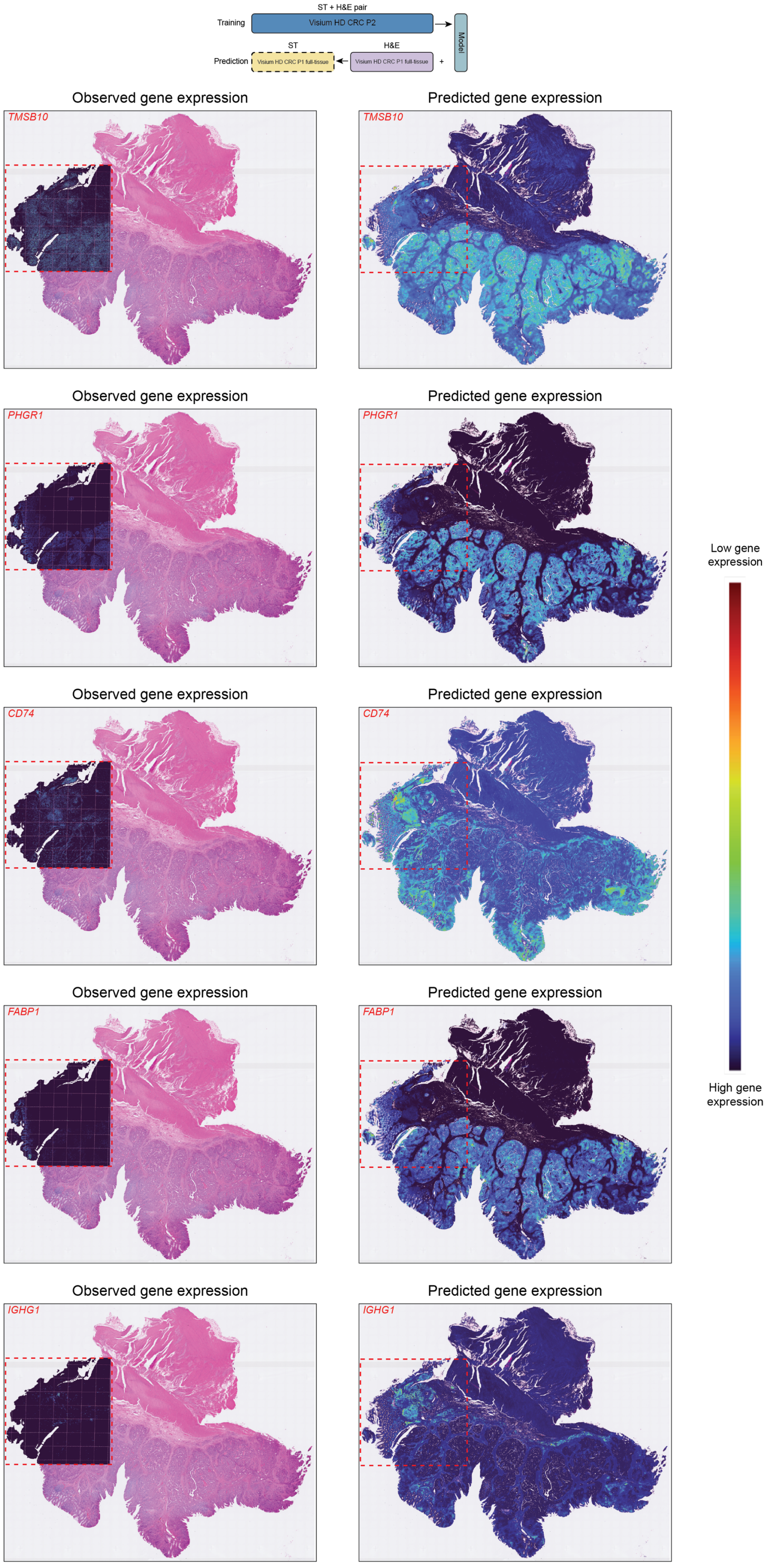

**Supplementary Fig. 6:** Additional examples of out-of-sample full-tissue prediction in CRC-P5. Pixel2Gene was trained on Visium HD CRC-P2 and applied to Visium HD CRC-P5. Full-tissue predicted expression for the same five genes shown in **Supplementary Fig. 4** is displayed alongside observed expression within the profiled region.

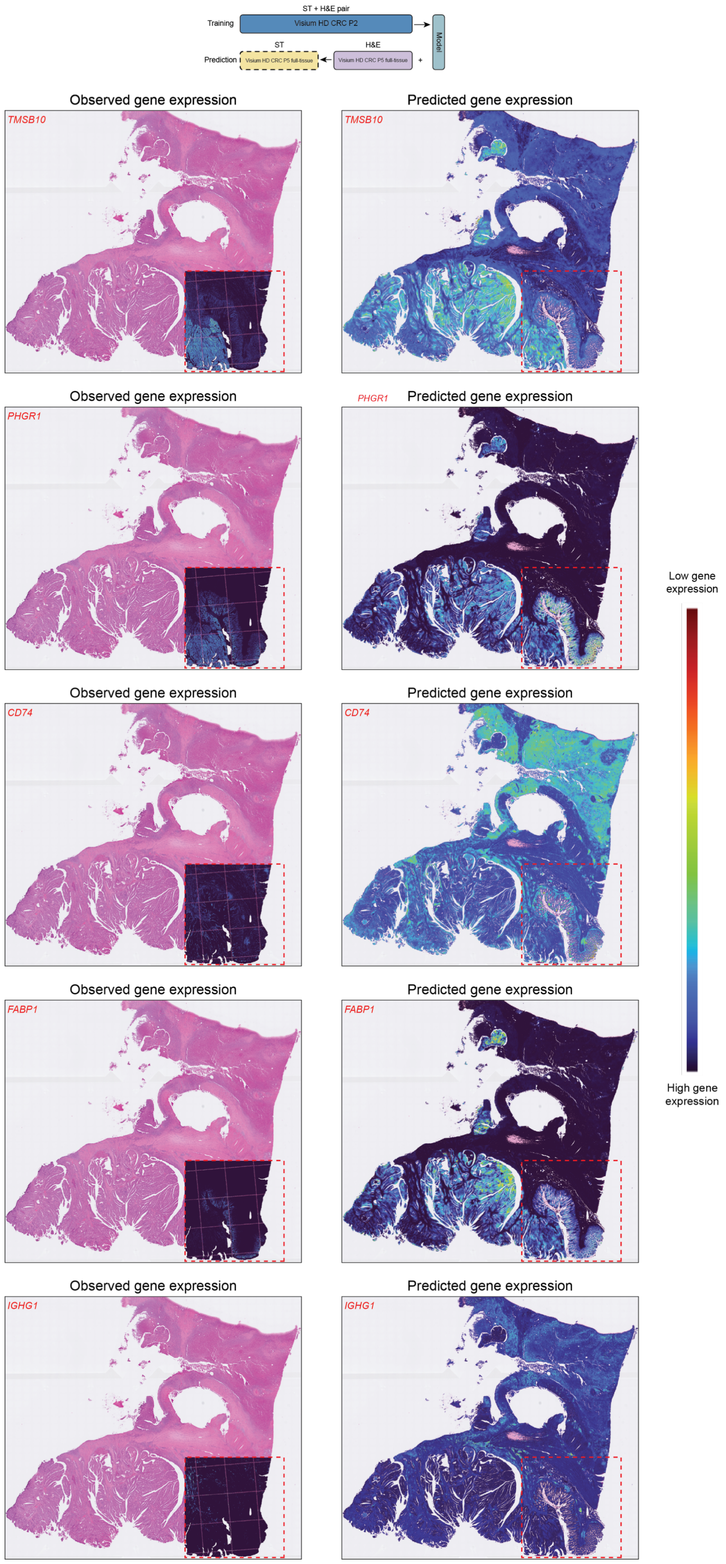

**Supplementary Fig. 7:** Additional examples of Pixel2Gene enhancement in Xenium CRC-P2. Pixel2Gene-enhanced spatial expression for five additional Xenium genes in CRC-P2. For each gene, observed Xenium expression is shown alongside Pixel2Gene-enhanced predictions.

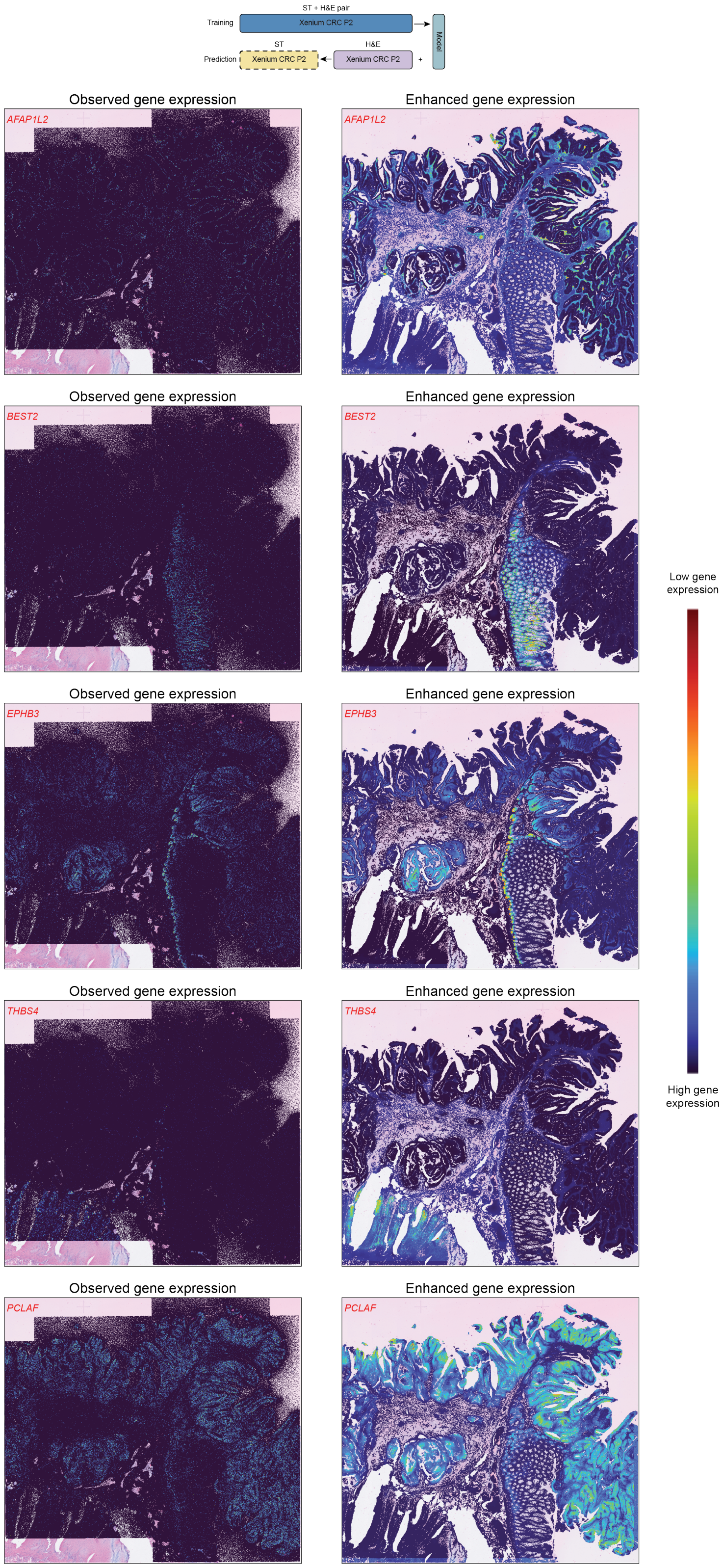

**Supplementary Fig. 8:** Out-of-sample Xenium prediction in CRC-P1 for additional genes. Pixel2Gene was trained on Xenium CRC-P2 and applied to Xenium sample CRC-P1. Predicted spatial expression for the same five genes shown in Supplementary Fig. 7 is displayed alongside observed Xenium expression.

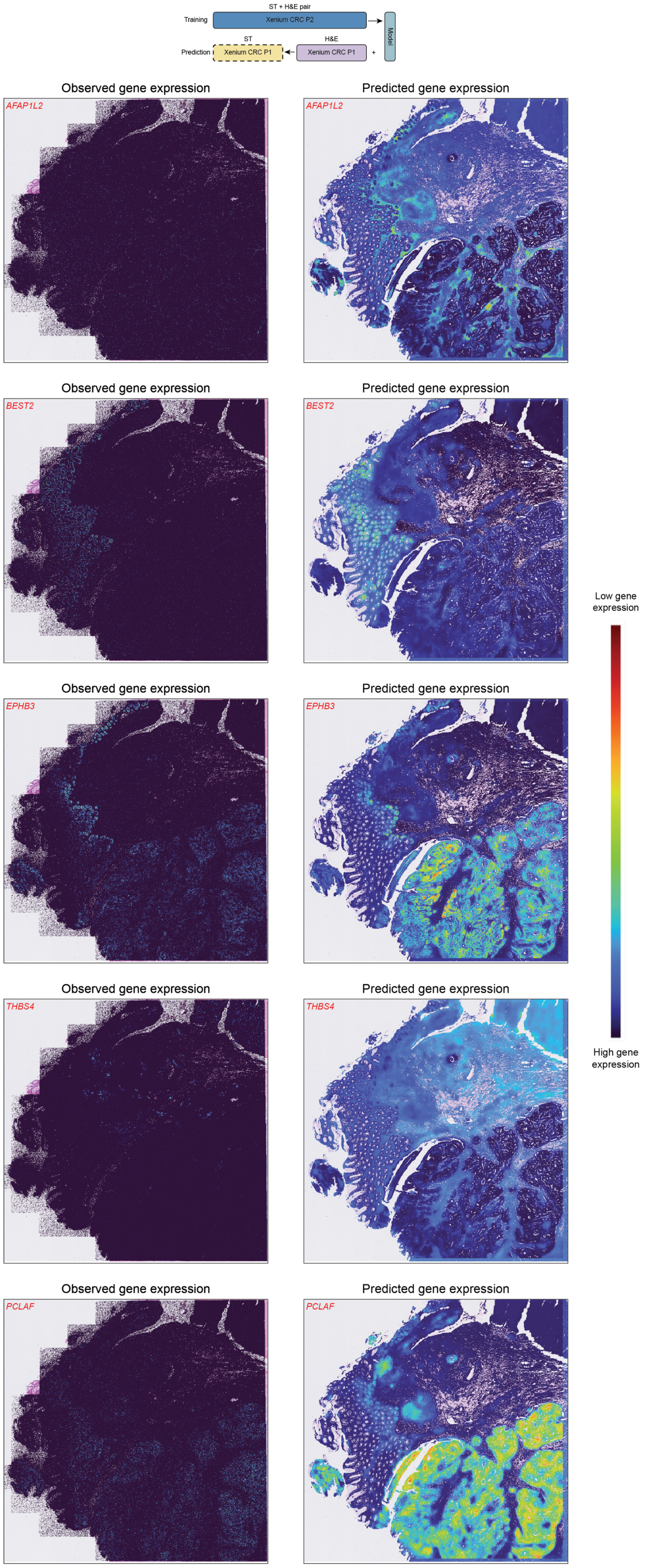

**Supplementary Fig. 9:** Out-of-sample Xenium prediction in CRC-P5 for additional genes. Pixel2Gene was trained on Xenium CRC-P2 and applied to Xenium sample CRC-P5. Predicted spatial expression for the same five genes shown in **Supplementary Fig. 7** is displayed alongside observed Xenium expression.

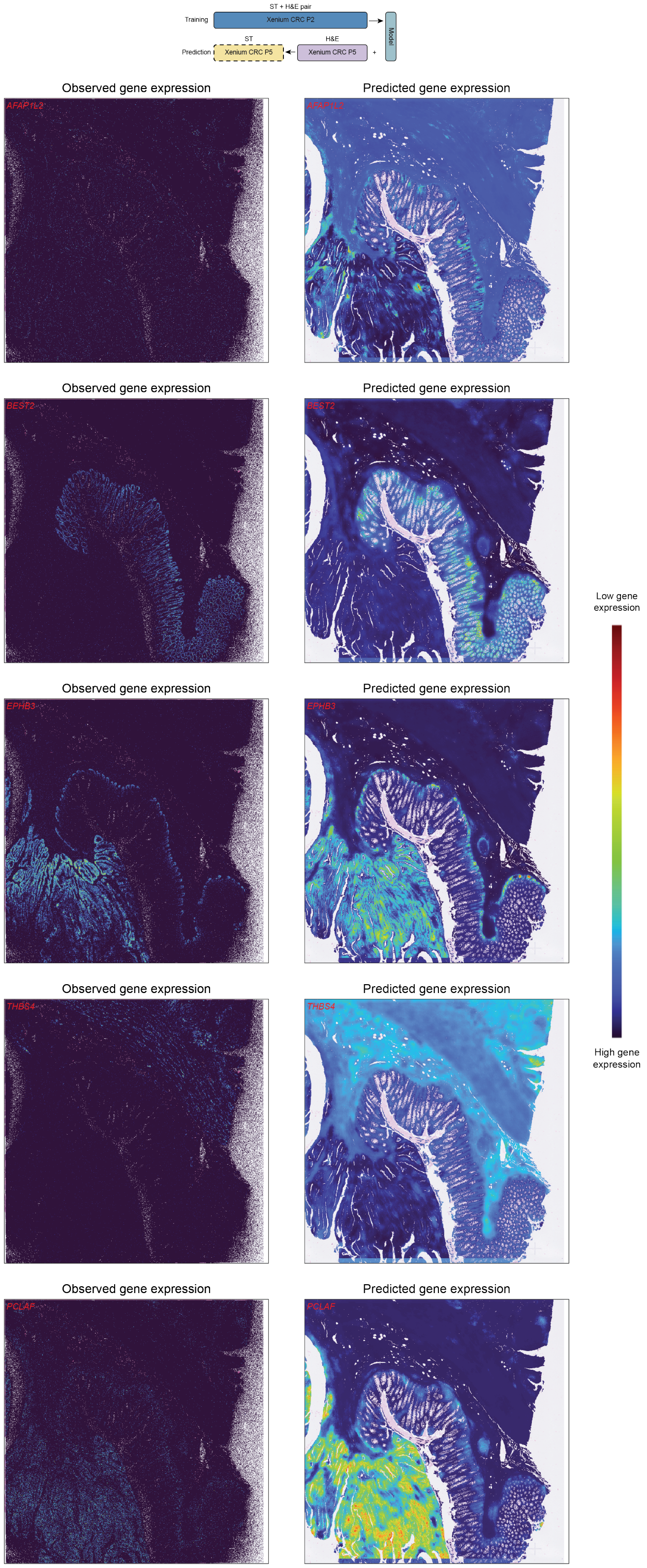

### References


1 Allen Institute for Brain Science. Reference Atlas :: Allen Brain Atlas: Mouse Brain. https://mouse.brain-map.org/static/atlas (2020).
